# Three dimensional nanoscopy of whole cells and tissues with *in situ* point spread function retrieval

**DOI:** 10.1101/727354

**Authors:** Fan Xu, Donghan Ma, Kathryn P. MacPherson, Sheng Liu, Ye Bu, Yu Wang, Cheng Bi, Tim Kwok, Peng Yin, Sarah Calve, Gary E. Landreth, Fang Huang

## Abstract

Single-molecule localization microscopy is a powerful tool in visualizing organelle structures, interactions, and protein functions in biological research. However, whole-cell and tissue specimens challenge the achievable resolution and depth of nanoscopy methods. As imaging depth increases, photons emitted by fluorescent probes, the sole source of molecular positions, were scattered and aberrated, resulting in image artifacts and rapidly deteriorating resolution. We propose a method to allow constructing the in situ 3D response of single emitters directly from single-molecule dataset and therefore allow pin-pointing single-molecule locations with limit-achieving precision and uncompromised fidelity through whole cells and tissues. This advancement expands the routine applicability of super-resolution imaging from selected cellular targets near coverslips to intra- and extra-cellular targets deep inside tissues. We demonstrate this across a range of cellular-tissue architectures from mitochondrial networks, microtubules, and nuclear pores in 2D and 3D cultures, amyloid-β plaques in mouse brains to developing cartilage in mouse forelimbs.

## INTRODUCTION

Fluorescence microscopy is a powerful and versatile tool in visualizing organelle structures, interactions, and protein functions in biological contexts. However, the smallest resolvable feature (i.e. the resolution) is restricted by the wave nature of light, typically ∼200 nm in the lateral plane and ∼500 nm in the axial dimension. This century-old barrier has restricted our understanding of cellular functions, interactions, and dynamics, particularly at the sub-micron to nanometer scale [1]. During the past decades, super-resolution fluorescence microscopy techniques, such as stimulated emission depletion (STED) microscopy [2–4], structured illumination microscopy (SIM) [5], and single-molecule localization microscopy (SMLM) [6–8] have overcome this fundamental limit offering up to 10-fold improvement in resolution and thus provided unprecedented opportunities to observe biological phenomena never before seen [9–15]. Specifically, SMLM (also known as PALM/STORM), as well as its three-dimensional (3D) counterpart [16–22], utilizes photo-switchable/convertible dyes or proteins to isolate emission patterns of individual molecules in different camera frames. By analyzing these information-carrying patterns, the positions of individual molecules can be pinpointed with a precision as low as 5 nm in three dimensions [9,14,15].

The core of 3D single-molecule super-resolution imaging can be summarized as encoding and decoding the single molecule positions in the form of their emission patterns (i.e. point spread functions (PSFs)). The encoding method, usually referred as PSF engineering, allows generating an emission pattern that changes, in an unambiguous and nondegenerate manner, with respect to the axial position of the single molecule [16–22]. The decoding method infers the location of a single molecule inside the biological specimen from the emission pattern detected on the camera. This inference process utilizes a wide range of well-established mathematical tools such as feature-based mapping [16,23], regression [24,25], and deep learning [26–28] to estimate the molecular position using a 3D PSF model, which describes the emission pattern of a single molecule with respect to its position within the specimen. It is, therefore, imperative to obtain an accurate model of the 3D PSF, which can reflect the complex biological and physical context constituting the path that the emitted photons travel through before being detected. Inaccurate models will give rise to imaging artifacts and significant deteriorations of the super-resolved 3D volume [29,30].

To account for instrument imperfections, both analytical and numerical methods have been developed to provide an accurate model that matches recorded emission patterns from fiduciary markers, in many cases using fluorescent beads on a coverslip surface [31–35]. In addition, by retrieving the PSF model from beads embedded within a layer of gel [36] or from fluorescent molecules coated on a latex microsphere [37], the effect of mismatched refractive indices between immersion oil and water-based imaging medium [38] can be characterized. However, none of these approaches takes the biologically heterogeneous and optically complex cell or tissue specimen into account. Therefore, these inherent complexities of individual specimens often render these *in vitro* calibrations inaccurate, especially when the intra- or extra-cellular targets are located away from the coverslip surface or inside thick tissues generating sample-induced aberrations [39]. These sample-induced aberrations result in emission patterns that change depending on the local and highly complex biological environment and establish a major challenge for the practical application of single-molecule super-resolution imaging in whole-cell and tissue specimens [40].

Here we propose a method to enable, for the first time, the construction of an *in situ* PSF response directly from the obtained single molecule dataset. Drawing inspiration from mathematical frameworks of expectation-maximization and *k*-means, the developed method allows us to eliminate the PSF mismatch and its resulting imprecision in localization induced by both instrument imperfections and the local biological context. The ability of retrieving accurate 3D PSF models *in situ* allows pin-pointing the positions of single molecules with uncompromised accuracy and precision, therefore, resolving intra- or extra-cellular structures within whole-cell and tissue specimens with high resolution and fidelity. We demonstrate its application across a range of cellular and tissue architectures from the nanoscale structures of mitochondrial networks, microtubules, and nuclear pore complexes throughout whole cells in 2D and 3D cultures, to amyloid β plaques in mouse brains and developing cartilage in embryonic mouse forelimbs.

## RESULTS

### Basic principle of *in situ* PSF retrieval (INSPR)

We start with a single molecule blinking dataset, routinely obtained in 3D SMLM experiments. In this dataset, each frame contains multiple isolated emission patterns originated from single molecules located in the cellular volume. Collectively, these single molecule patterns can be regarded as random observations of emission patterns at various axial positions from the 3D PSF that we want to retrieve. Thousands to millions of such emission patterns are detected in each single molecule dataset. If correctly combined, these single molecule detections will provide an adequate reconstruction of their underlying 3D response function (i.e. a 3D point spread function). The key that links these acquired single molecule patterns to our desired 3D response is the molecular positions of the single emitters, in particular, their axial positions. This key, however, is missing.

We draw inspiration from expectation-maximization [41] and *k*-means frameworks [42,43] to retrieve the 3D PSF response in the presence of unobserved latent parameters – the axial and lateral positions of single molecules (Figure 1, Figure S1A). This *in situ* PSF retrieval algorithm (referred as ‘INSPR’ hereafter, for simplicity) iteratively uses two separate steps, namely assignment and update, to build an *in situ* PSF model from a collection of single molecule patterns. Pupil function, representing the wave field at the pupil plane of the microscope, is used to succinctly describe the 3D PSF response at arbitrary axial positions. In brief, INSPR starts with an ideal PSF (i.e. with a constant pupil) and then assigns each detected single molecule pattern to a temporary axial position through cross correlation with this ideal template. These axially assigned single molecule patterns are subsequently grouped, aligned, and averaged to form a 3D PSF stack, which provides a new pupil estimation (an ‘update’ to the previous pupil) through phase retrieval. This new pupil is then used in the next assignment step to generate an updated template. This process iterates until good agreement between the detected PSFs and the retrieved model is reached (see **Supplementary Notes** for detailed description). We find the algorithm converges relatively fast within 6–10 iterations (convergence criteria: phase difference <0.02 *λ*), depending on the PSF complexity.

**Figure 1.**
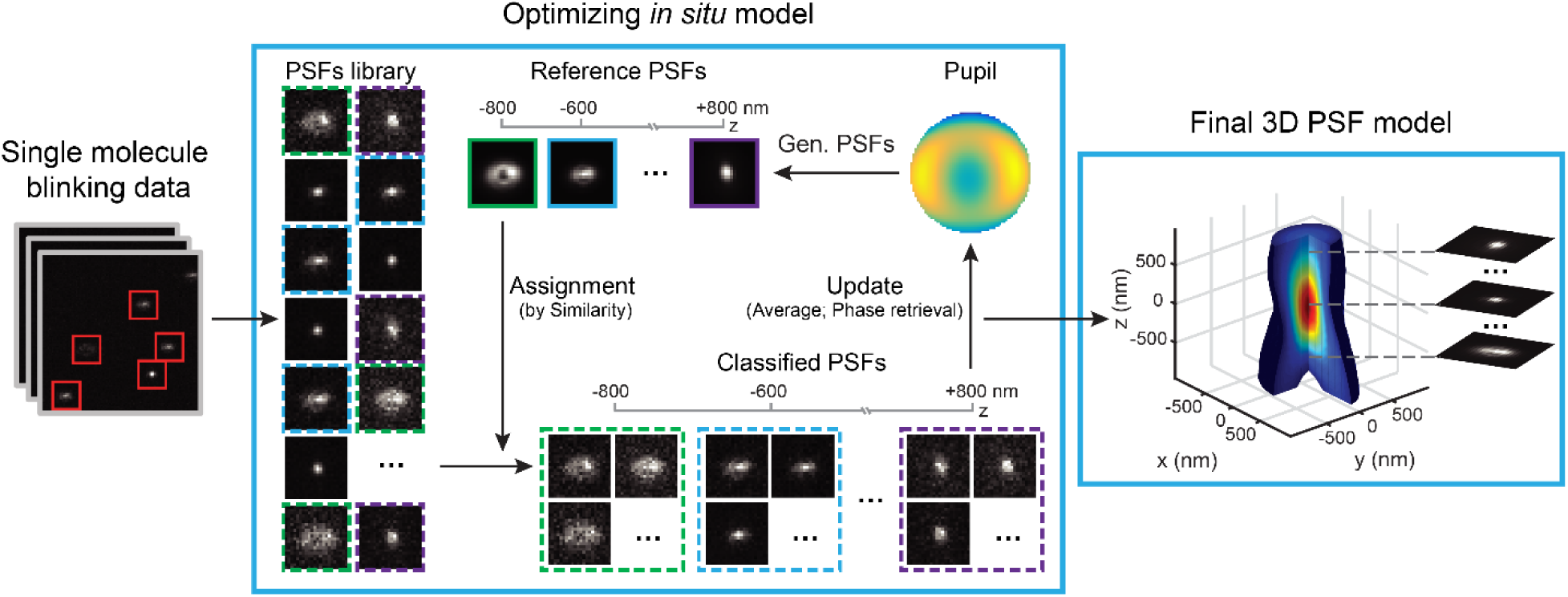
Concept of *in situ* PSF retrieval (INSPR). After single molecule blinking data (left panel) is acquired, a PSFs library is obtained. Starting with a constant pupil function, INSPR assigns each detected PSF to a temporary axial position according to its similarity with this ideal template (assignment step, center panel). These axially-assigned PSFs are subsequently grouped, aligned, and averaged to form a 3D PSF stack. This 3D PSF stack is then used to provide a new pupil estimation through phase retrieval (update step, center panel). This new pupil is used in the next assignment step to generate an updated template. This process iterates until good agreement between the detected PSFs and the retrieved model is reached, and then a final 3D PSF model is obtained (right panel). Gen. PSFs stands for generating PSFs.

To build a unique *in situ* PSF model, the 3D single-molecule imaging modality must avoid degeneracies. Degeneracy appears when more than one state, a set of variables being estimated, lead to the same observation. One such state of INSPR is the wavefront shape, which describes the aberration introduced by the imaging system and specimen and can be decomposed by a series of Zernike polynomials [44]. Degeneracy arises when two different wavefront shapes (i.e. two different states) generate the same single molecule emission pattern (i.e. the observation). For example, positive and negative vertical astigmatism aberrations will generate identical PSF patterns at opposite axial positions (Figure S1C), making them impossible to be classified in the assignment step. We break up these degeneracies by using a biplane configuration (Figures S1F and S1G), where a pair of emission patterns from the same single molecule is detected at two axially separated planes (Figure S1D). By registering this pair of PSFs in the assignment step, we can retrieve the *in situ* 3D PSF without ambiguity (**Supplementary Notes**).

To pin-point single molecule positions with high precision and minimum bias, we combine INSPR with maximum likelihood estimator (MLE) that incorporates sCMOS (scientific complementary metal-oxide-semiconductor) camera-specific pixel-dependent noise model [45] to allow applications that rely on fast acquisition speed (e.g. in live-cell imaging) and large field of view (e.g. in high-throughput studies) offered by the CMOS sensor. To enable unaltered statistical properties of isolated single molecule emission patterns and their associated photon-counting and pixel-dependent noise, INSPR generates a channel-specific *in situ* PSF for each detection plane separately (Figure S1B, **Supplementary Notes**). This approach avoids transforming and interpolating the single molecule dataset and, therefore, allows reducing imaging artifacts and localization imprecision when using multiple detection planes, as the statistical properties of the raw detected camera counts will be lost during transformation (Figures S2E and S2F).

### Performance quantification of *in situ* PSF retrieval with INSPR

Next, we test the accuracy of INSPR by retrieving a known wavefront distortion from a library of single molecule emission patterns simulated randomly within an axial range of ±800 nm (Figure 2A, Movie S1). The known wavefront shape consists of 21 Zernike modes (Wyant order, from vertical astigmatism to tertiary spherical aberration) with their amplitudes randomly sampled from –1 to +1 (unit: *λ*/2π). We find INSPR has successfully retrieved the *in situ* pupil with a phase error of 15±6 m*λ* (30 trials) (Figure 2B, Movie S1), and a Zernike amplitude error of 11±4 m*λ* (30 trials) for the total 21 modes (Figure 2C, Movie S1). The resulting PSFs generated by the *in situ* pupil show high degree of similarity with those generated by the known pupil (Figure 2D). The performance is further tested through retrieving a previously estimated wavefront distortion at various imaging depths above the coverslip, up to ∼45 μm [30] (Figures S2A and S2B). We find that INSPR enables retrieval of these *in situ* PSFs that vary significantly throughout the depths (0, 6.7, 14.35, 27.55, and 45.4 μm). While these highly-distorted wavefronts make decomposition into Zernike modes unreliable due to phase wrapping, we find the normalized cross correlation coefficients between the retrieved *in situ* PSF and the known response can be consistently maintained above 0.95 at each tested depth within a relatively large axial range (±800 nm) (Figure S2A).

**Figure 2.**
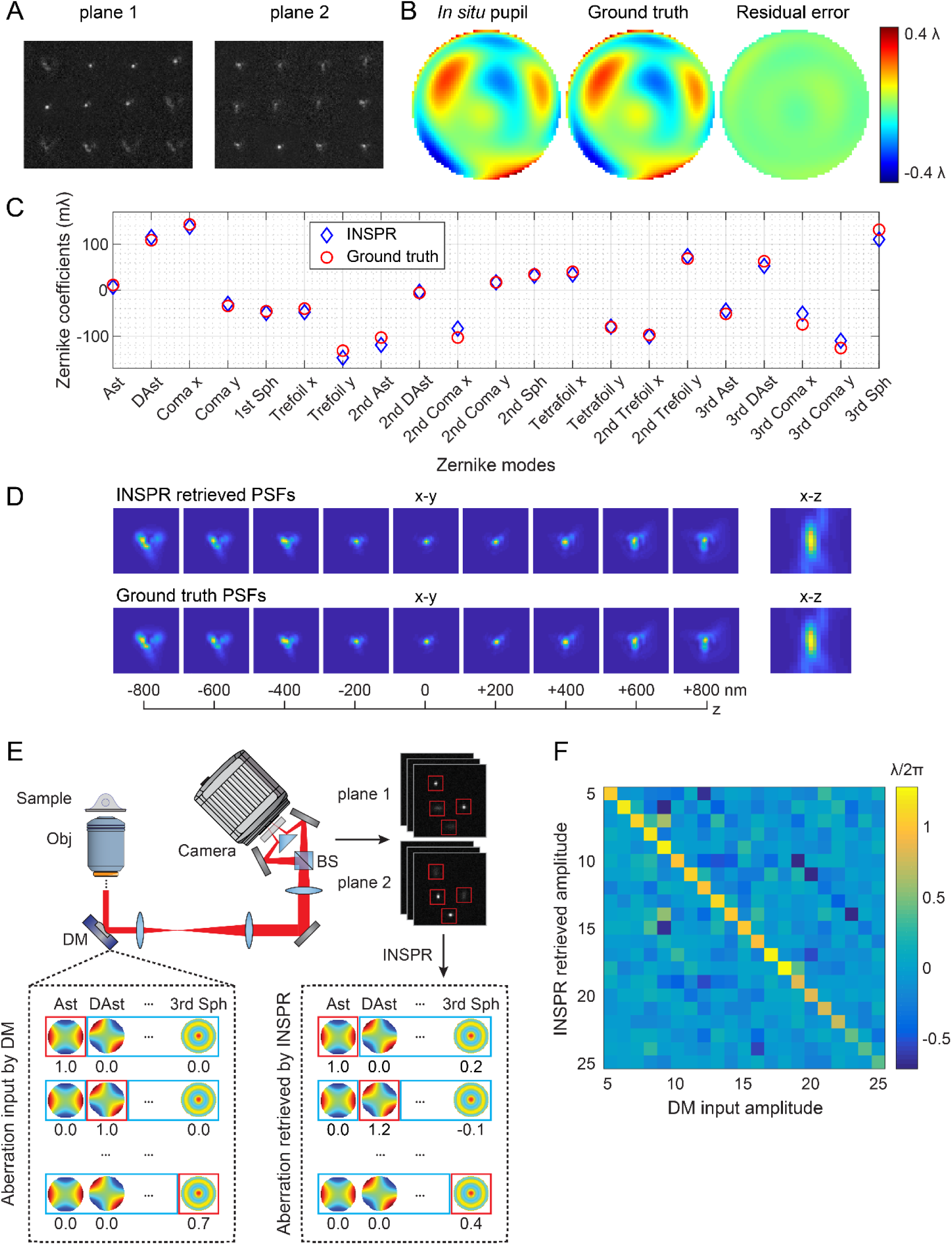
Performance quantification of *in situ* PSF retrieval with INSPR. (A) Simulated biplane single molecule emission patterns located randomly over an axial range from –800 to +800 nm with a known wavefront distortion. An animated demonstration of the total 30 trials is shown in Movie S1. (B) Phase of the *in situ* pupil retrieved by INSPR (left), ground truth (middle), and the residual error (right, 15.6 m*λ*). (C) Amplitudes of 21 Zernike modes (Wyant order, from vertical astigmatism to tertiary spherical aberration) decomposed from the *in situ* pupil retrieved by INSPR (blue diamonds), with a root-mean-square error (RMSE) of 11.1 m*λ* for the total 21 modes compared with ground truth (red circles). (D) x-y and x-z views of the INSPR retrieved PSFs (top row), showing high degree of similarity with those generated by the known pupil function (bottom row). (E) Schematic for testing the performance of INSPR in a cellular dataset of immuno-labeled TOM20, a mitochondria marker, using DNA-PAINT, where a deformable mirror is inserted in the pupil plane of the microscope to introduce controllable wavefront distortions. Single molecule emission patterns distorted by the input Zernike-based aberrations are acquired, and then fed into INSPR to retrieve the *in situ* pupil and its corresponding aberrations. (F) Heat map showing the relationship between the deformable mirror input and INSPR retrieved amplitudes of 21 Zernike modes. Compared with the amplitudes of the aberrations input by the deformable mirror along the diagonal elements, INSPR results in 8% estimation error for the first 18 Zernike modes, and 41% error for the last three tertiary aberration modes. An animated process to generate this heat map is shown in Movie S2. Ast: vertical astigmatism; DAst: diagonal astigmatism; Sph: spherical aberration; Obj: objective lens; DM: deformable mirror; BS: 50/50 non-polarizing beam splitter cube.

By inserting a deformable mirror in the pupil plane (also known as Fourier plane) of the microscope, we can introduce controllable wavefront distortions mimicking situations when imaging through whole-cell and tissue specimens (Figure 2E, Figure S1F). We calibrated the deformable mirror (**Supplementary Notes**) to introduce individual Zernike-based aberration modes ranging from commonly experienced astigmatism and coma to high-order modes such as tertiary spherical aberration. We acquired single molecule blinking datasets in COS-7 cells by visualizing immuno-labeled mitochondrial marker TOM20 through DNA-PAINT (DNA point accumulation for imaging in nanoscale topography) [46–49]. The introduced aberrations distorted the single molecule emission patterns detected on the camera, which were then fed into INSPR to retrieve the *in situ* PSF and its corresponding pupil function. By comparing the aberration amplitudes induced by the deformable mirror with those retrieved by INSPR (Figure 2F, Movie S2), we find that INSPR provides accurate estimations for the first 18 Zernike modes (Wyant order, 8% error compared to the phase retrieval result using beads *in vitro*, Figure S2D, **Supplementary Notes**) with a performance decrease in the last three tertiary aberration modes (41% error, **Supplementary Notes**). In agreement with our previous results, this result demonstrates the capability of INSPR in retrieving distorted *in situ* PSFs directly from single molecule data obtained in cellular contexts.

INSPR depends on the stochastic switching of single molecules to reconstruct the underlying PSF. Consequently, its convergence rate, i.e. the number of emission patterns needed for a stable reconstruction (convergence criteria: Zernike amplitude error <0.02 *λ*), depends on the signal to background ratio (SBR) of the detected emitters (Figure S2C). We find that, in high SBR cases, a condition usually encountered when using specific labeling methods such as DNA-PAINT or bright organic probes such as Alexa Fluor 647, INSPR requires less than 300 emission patterns to converge. In contrast, INSPR requires more than 2100 emission patterns in low SBR cases, a common condition for live-cell single-molecule experiments with fluorescent proteins such as mEos3.2 [50]. For fixed-cell imaging, INSPR requires a relatively small number of emission patterns to converge when compared to the total obtained single molecule patterns. However, for live-cell imaging, the required number of emission patterns might limit the temporal resolution of INSPR when rapid temporal variation of wavefront distortion is sought.

### Three-dimensional super-resolution imaging of whole cells with INSPR

INSPR enables us to measure and compensate sample-induced distortions within the actual imaging volume, as well as capturing its evolution throughout a thick specimen. The conventional method relies on the PSF model generated by using fluorescent beads attached on a coverslip surface and uses it to localize single molecule emission events in all depths of a biological specimen (hereafter referred as the *in vitro* approach). To the contrary, INSPR builds a specific *in situ* PSF model from the acquired single molecule dataset in each optical section and uses this model to localize the emission events in the corresponding section with high accuracy.

As a demonstration, we imaged immuno-labeled TOM20 protein, a protein complex located in the outer membrane of mitochondria, in COS-7 cells (Figure 3, Figure S3, Movie S3). To investigate the feasibility of INSPR when imaging above the coverslip surface, we created a 9-µm-thick sample cavity filled with the water-based imaging medium between two coverslips, with the immuno-labeled cells on the upper coverslip. A volume of ∼30 µm × 30 µm × 2.5 µm within a COS-7 cell was imaged. We find that the interconnected mitochondrial network can be clearly resolved with high resolution by using INSPR (Figure 3A), where the horizontal and vertical cross sections reveal individual TOM20 clusters distinctly (Figures 3B and 3C). Examining reconstructions of the same field of view from both INSPR and the conventional *in vitro* approach, we find INSPR is capable of resolving the surface contour of each organelle with high resolution in all three dimensions (Movie S3) whereas reconstructions using the *in vitro* approach exhibit both artifacts and decreased resolution in these examined cross sections (Figures 3D–3K, Figure S3B). Intensity profiles of 25 typical outer membrane contours (positions shown in Figure S3A) demonstrate a consistent improvement in resolution achieved by INSPR in comparison to the *in vitro* approach (Figure 3L, Figure S3E) with a mean precision of ∼8 nm in lateral and ∼21 nm in axial dimensions as estimated by the Cramér-Rao lower bound (CRLB) (Table S1, **Supplementary Notes**).

**Figure 3.**
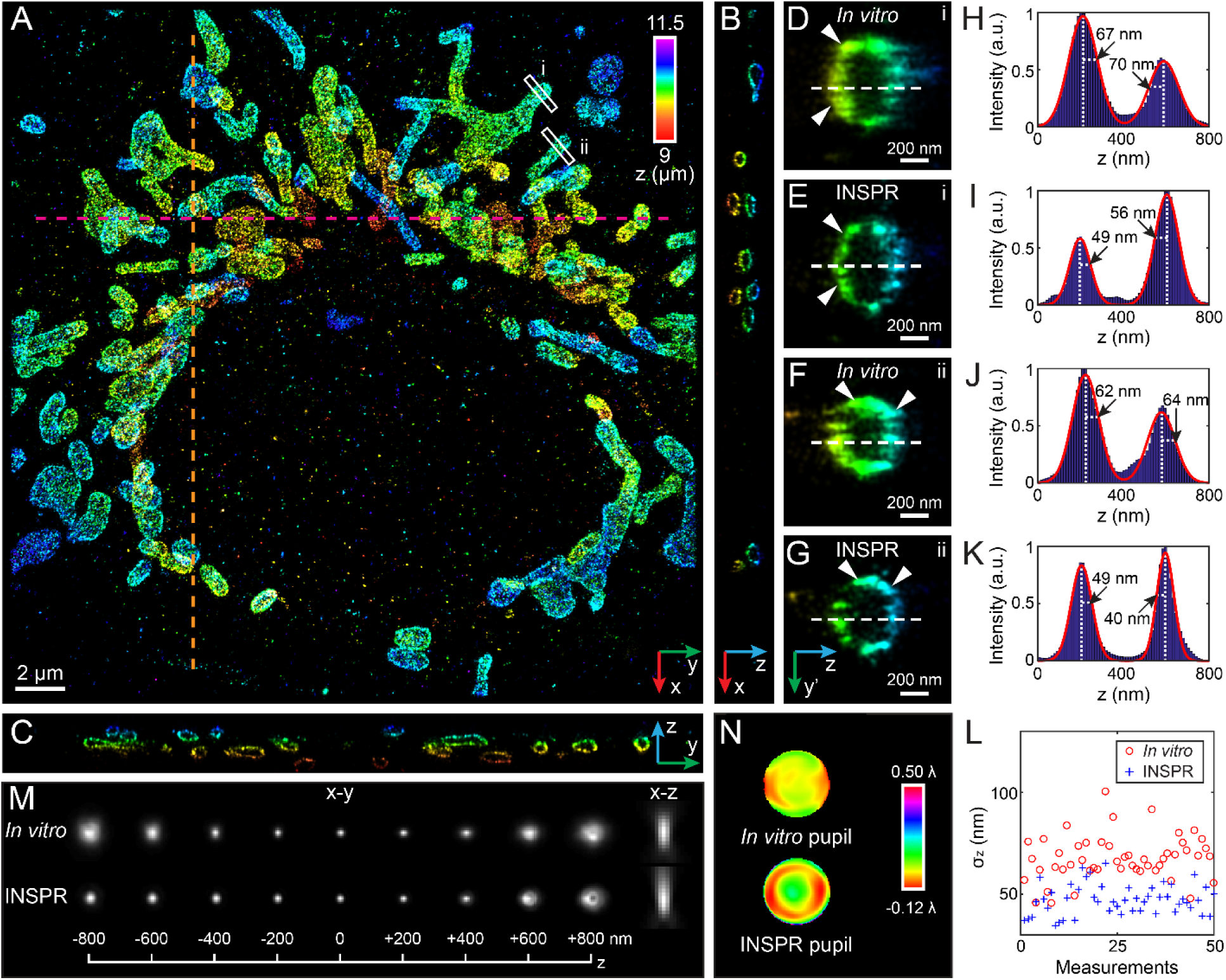
Comparison of 3D reconstructions of the mitochondrial network labeled with TOM20 and the retrieved 3D PSFs using INSPR and the *in vitro* method. (A) x-y overview of the mitochondrial network beyond a 9-µm-thick sample cavity filled with the water-based imaging medium from the bottom coverslip. The interconnected network is clearly resolved by INSPR with high resolution. The depth information is color-coded according to the color bar. An animated 3D reconstruction is shown in Movie S3. (B and C) x-z and y-z slices along the orange and magenta dashed lines in (A), respectively, where individual TOM20 clusters are revealed distinctly. The thickness of these axial slices along the third dimension is 200 nm. (D–K) Enlarged y’-z views of the outer membrane structures in areas i (D and E) and ii (F and G) as indicated by the white-boxed regions in (A), and their intensity profiles along the z direction (H–K). The surface contour of each organelle is resolved by INSPR with high resolution in all three dimensions (E and G), while reconstructions using the *in vitro* approach exhibit artifacts (D and F, as highlighted by arrowheads). Intensity profiles of these outer membrane contours show higher resolution achieved by INSPR (I and K) compared with the *in vitro* approach (H and J). Here the orientation of the cross section is rotated to allow projection of the 3D membrane bounded structures to the 2D image. (L) Distribution of σ_z_ obtained from the intensity profiles of 25 typical outer membranes in (A). INSPR results are marked with blue plus signs and the *in vitro* results are marked with red circles. (M) x-y and x-z views of the PSFs generated by the INSPR model of the deepest section above the bottom coverslip (bottom row) and those generated by the *in vitro* model (top row). The axially stretched PSFs retrieved by INSPR show the existence of sample-induced aberrations when imaging deep into the cell, which cannot be observed by imaging fluorescent beads on a coverslip surface. (N) Phase of the INSPR pupil and the *in vitro* pupil corresponding to PSFs as shown in (M). The INSPR pupil shows sample-induced aberrations together with instrument imperfections, while the *in vitro* pupil can characterize instrument imperfections but fails to reflect sample-induced aberrations.

To further explain this difference, we compare the *in situ* PSF models retrieved by INSPR with the *in vitro* one (Figures 3M and 3N, Figures S3C and S3D). As the imaging depth increases, the amount of sample-induced aberrations from optical sections does so accordingly. The *in situ* pupils retrieved by INSPR (Figure 3N, Figure S3C), together with their decomposed Zernike amplitudes (Figure S3D), faithfully reflect this phenomenon as a progressive increase of major aberrations such as spherical and coma with the increasing depths. This is also shown in the axially stretched PSFs retrieved by INSPR (Figure 3M, Figure S3C). In contrast, the PSF retrieved from fluorescent beads successfully characterizes instrument imperfections but fails to take into account sample-induced aberrations and their depth-dependent variations due to its *in vitro* nature.

Given the ability of retrieving an *in situ* 3D response and therefore pin-pointing single molecules with high accuracy, we tested INSPR by reconstructing nucleoporin Nup98 immuno-labeled with Alexa Fluor 647 in COS-7 cells (Figure 4, Figure S4, Movie S4), which localizes near the center channel of the nuclear pore complex (NPC) [51,52]. We first reconstructed a super-resolution 3D volume of Nup98 within a relatively small depth of 3.3 µm and found individual ring-like structures covered the bottom surface of the nucleus envelope displaying slight invaginations and undulations (Figures 4A–4D). We then reconstructed Nup98 on the entire nuclear envelope with a total thickness of 6.4 µm (Figures 4E–4L), and found that not only the individual pores were distinctly resolved throughout the entire envelop, but also their ultra-structures were resolved at both bottom and top surfaces of the nucleus (Figures 4F and 4G, Movie S4). Quantifying these resolved nanoscale structures of Nup98, we find the average diameters are 60±9 nm and 57±11 nm (Figure S4E, 40 measurements for each sample, profile positions shown in Figures S4A and S4C), which is consistent with being localized to the NPC channel walls and its labeling using IgG antibody molecules. Quantitatively, lateral profiles of single boundaries (i.e. ring thickness) of the observed structures result in σ_lateral_ of 14±3 nm and 11±3 nm (Figure S4F, 80 measurements for each sample, profile positions shown in Figures S4A and S4C), while we observe σ_z_ of the top envelop surface (46±12 nm) is similar to that of the bottom surface (48±9 nm and 36±11 nm) (Figure S4G, 20 measurements for the bottom surface of the 3.3-µm-thick sample and 10 measurements for each surface of the 6.4-µm-thick sample, profile positions shown in Figures S4B and S4D). The localization precision estimated by CRLB achieves ∼10 nm in lateral and ∼35 nm in axial dimensions (Table S1).

**Figure 4.**
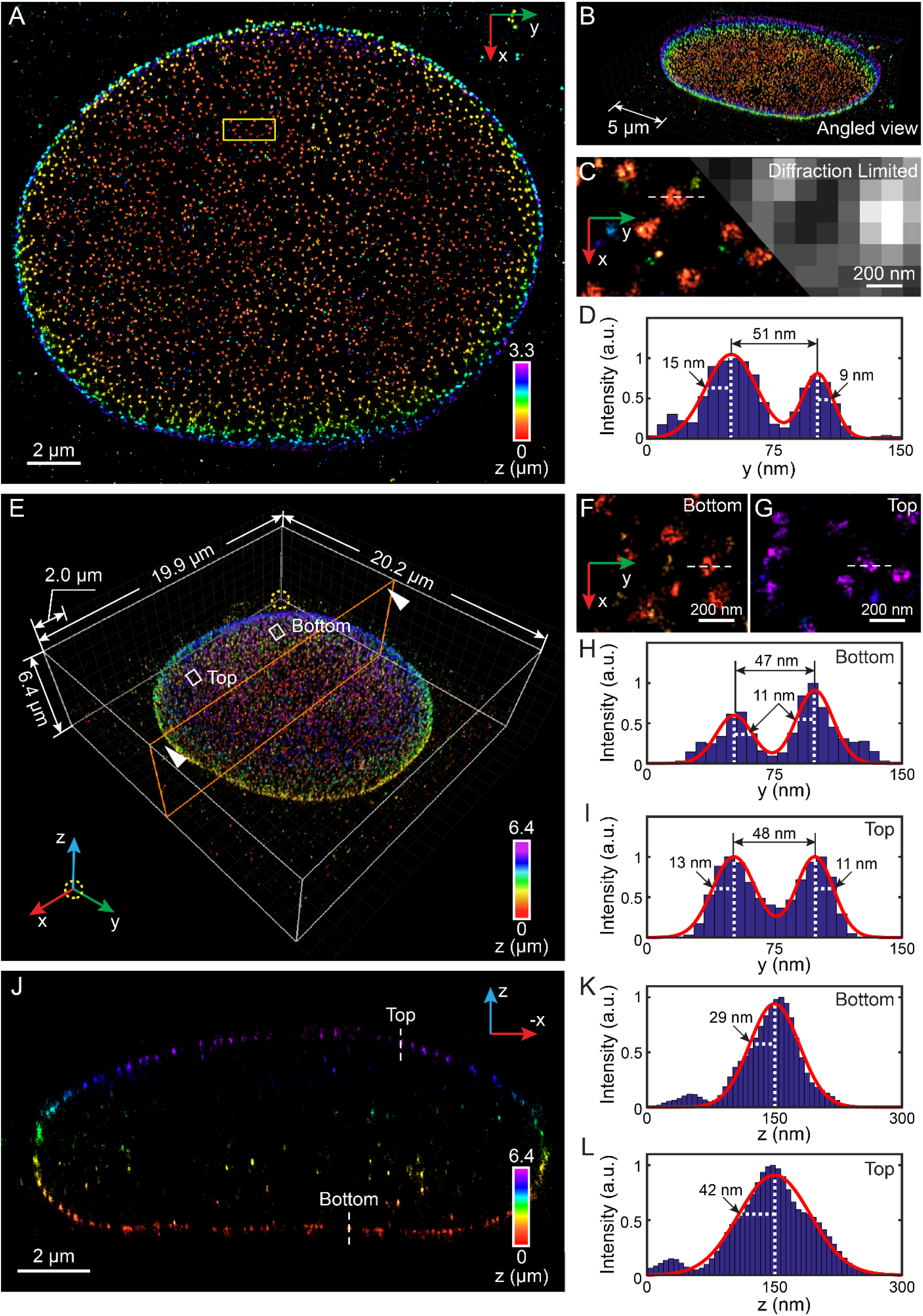
3D super-resolution reconstruction of nuclear pore complex protein Nup98 on the nuclear envelope. (A) x-y overview of a 3.3-µm-thick section of the nucleus, where individual ring-like structures cover the bottom surface of the nucleus envelope with slight invaginations and undulations. (B) Angled view of (A). (C) Sub-region as indicated by the yellow-boxed region in (A) showing the nanoscale structure of Nup98 (left), which are blurred and not resolvable in conventional diffraction-limited microscopy (right). (D) Intensity profile along the white dashed line in (C). The measured diameter of this Nup98 structure is 51 nm, while σ_y_ obtained from the left and right boundaries equals to 15 nm and 9 nm, respectively. (E) Nup98 on the entire nuclear envelope rendered in 3D, with a total thickness of 6.4 μm. The axial localization of the nucleus is color coded in z. An animated 3D reconstruction is shown in Movie S4. (F and G) Sub-regions as indicated by the white-boxed regions in (E) showing enlarged x-y views of nuclear pores, where the ultra-structures are resolved at both bottom (F) and top (G) surfaces of the nucleus. (H and I) Intensity profiles along the white dashed lines in (F) and (G). The diameters of the Nup98 structures are 47 nm and 48 nm at the bottom (H) and top (I) surface, respectively. The mean of σ_y_ is 11.5 nm, similar to the measured value in (D). (J) x-z cross section of the orange plane in (E) showing high resolution in the axial direction. The thickness of this cross section is 500 nm. (K and L) Intensity profiles along the white dashed lines in (J). σ_z_ equals to 29 nm for the measured structure at the bottom surface (K), and 42 nm for that at the top surface (L).

### Resolving amyloid β fibrils in mouse brain sections

Due to the inhomogeneous refractive indices of intra- and extra-cellular constituents, tissue specimens challenge optical imaging in its achievable resolution and depth. Due to the density of these materials packed within a small inter-connected volume, photons emitted by single fluorescent probes, the sole source of the molecular position, are aberrated and scattered when traveling through the tissue specimen, resulting in distorted emission patterns. From the collection of these distorted single molecule emission patterns, INSPR can retrieve the underlying *in situ* PSF and pin-point these emission patterns within the tissue volume with high precision and accuracy. Here we imaged extracellular deposits of amyloid β (Aβ) in brain sections from an 8-month-old murine model of Alzheimer’s disease (5xFAD), which was routinely used to assess biological responses associated with Aβ accumulation (Figure 5, Figures S5A–S5D, Movies S5 and S6). In 8-month-old aging animals, the increasing amyloid burden is associated with cognitive deficits, gliosis, and neuroinflammation [53], while the pathological mechanism remains unclear. Quantification of Aβ based on conventional microscopy methods can result in contradictory findings [54] due to their insufficient resolutions. By using INSPR, we reconstructed various Aβ plaques with depths up to 16 µm in 30-µm-thick brain slices (Figure 5, Figures S5A–S5D). In a reconstructed super-resolution volume with a relatively low density of fibrils (Figures 5A–5J), we find that the distinct arrangement of fibrils in the plaque center can be resolved by using INSPR, and the 3D details of individual fibrils within the intercrossing fibril networks can be explicitly visualized and traced as demonstrated in cross sections of both x-y and y-z planes (Figures 5C–5J, Movie S5). In another reconstructed volume with a higher density of fibrils (Figures 5K–5P), we observe two distinct and axially separated layers of an Aβ plaque, each of which contains a network that is highly intercrossed with distinctly resolved fibrils (Figures 5M–5P, Movie S6). Measuring the 3D cross section of plaque forming fibrils resolved using INSPR, we obtain lateral profile widths (quantified with full width half maximum, FWHM) of 53±9 nm and 55±11 nm (Figure 5Q, 40 measurements in each volume, profile positions shown in Figures S5A and S5C) and axial widths of 112±31 nm and 118±21 nm (Figure 5R, 40 measurements in each volume) with an estimated localization precision of ∼9 nm in lateral and ∼33 nm in axial dimensions (by CRLB, Table S1). These results demonstrate the ability of INSPR to capture and discern individual fibrils within Aβ plaques of a 30-μm-thick AD brain slice while maintaining the high-resolution capability of single-molecule super-resolution techniques throughout the depth, which could further allow investigating interactions of Aβ species and tau aggregates within neurons, adjacent astrocytes, and microglial cells.

**Figure 5.**
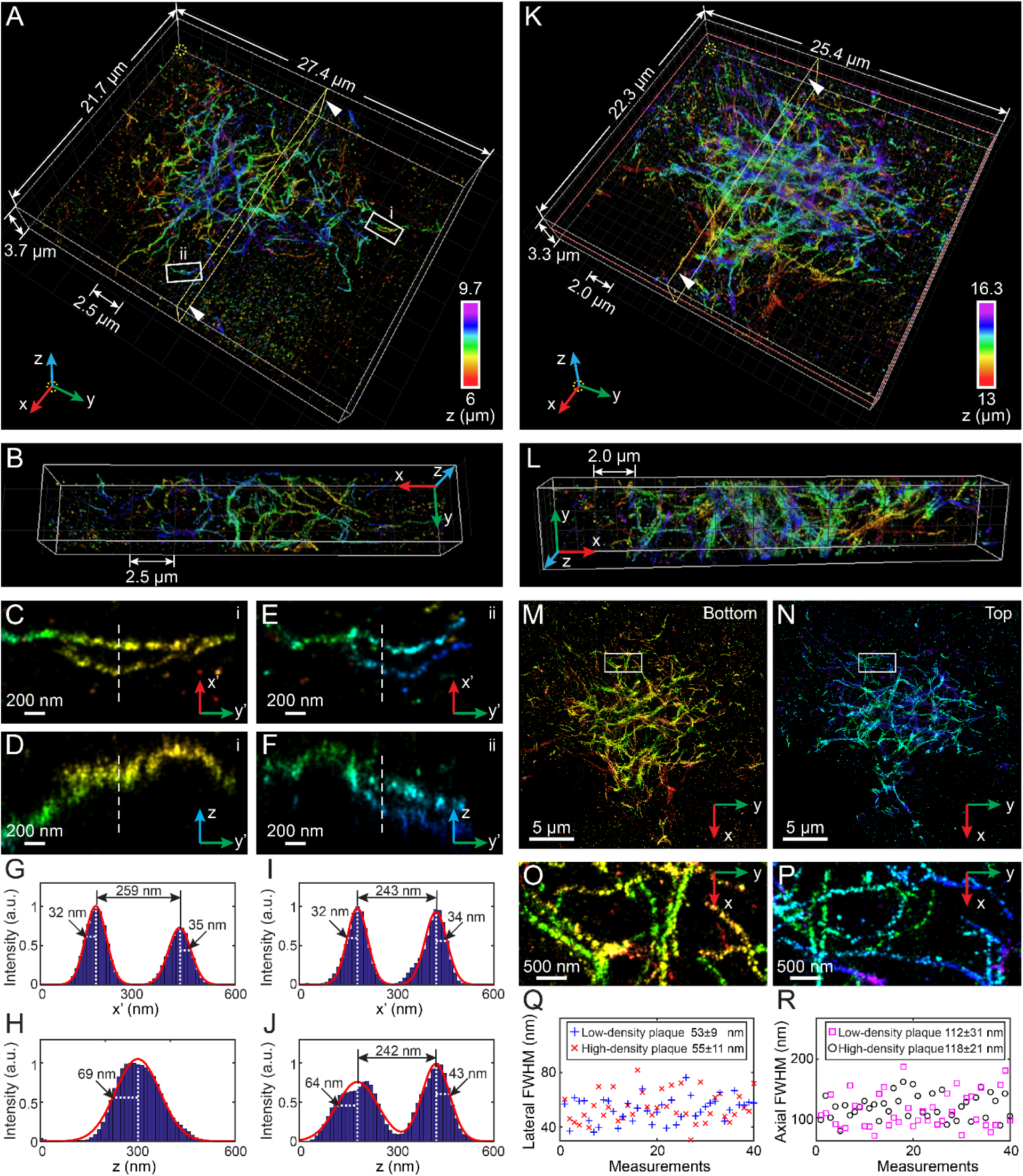
3D super-resolution reconstruction of amyloid β (Aβ) plaques in 30-μm-thick brain sections of mouse models of Alzheimer’s disease. (A) Overview of a reconstructed super-resolution volume of an Aβ plaque with a relatively low density of fibrils at a depth of 6 µm with a thickness of 3.7 µm. An animated 3D reconstruction is shown in Movie S5. (B) Cross section of the yellow plane in (A) showing the distinct arrangement of fibrils in the plaque center. (C–F) Zoomed in y’-x’ and y’-z views of two typical fibrils as indicated by the white-boxed regions i (C and D) and ii (E and F) in (A). Here the orientation of the cross section is rotated to allow projection of the 3D fibrils to the 2D image. (G–J) Intensity profiles along the white dashed lines in (C–F). (K) Overview of a reconstructed super-resolution volume of an Aβ plaque with a relatively high density of fibrils at a depth of 13 µm with a thickness of 3.3 µm, where two distinct and axially separated layers of an Aβ plaque are observed. An animated 3D reconstruction is shown in Movie S6. (L) Cross section of the yellow plane in (K) showing the interconnections among these dense fibrils. (M and N) x-y views of the bottom (M) and top (N) half of the plaque as divided by the orange plane in (K), each of which contains a network with distinctly resolved Aβ fibrils. (O and P) Enlarged x-y views of the areas as indicated by the white-boxed regions in (M) and (N) showing the different arrangement of fibrils in two layers. (Q) Distribution of lateral profile widths (quantified by FWHM) measured from 40 typical fibrils in the x-y plane in each plaque. The lateral FWHM is 53±9 nm for the low-density plaque (blue plus signs), and 55±11 nm for the high-density plaque (red crosses). (R) Distribution of axial FWHM measured from 40 typical fibrils in the x-z plane in each plaque. The axial FWHM is 112±31 nm for the low-density plaque (magenta squares), and 118±21 nm for the high-density plaque (black circles).

### Revealing elastic fiber structures in developing cartilage

As a final demonstration of INSPR in resolving complex tissue architectures, we imaged developing cartilage in the humerus of E14.5 mouse embryos (Figure 6, Figures S5E and S5F, Movie S7). To increase the signal to background ratio and focus on the extracellular matrix (ECM), the tissue was decellularized with 0.05% SDS and stained with wheat germ agglutinin (WGA) conjugated to Alexa Fluor 647. WGA is a lectin that binds to n-acetyl glucosamine and sialic acid, enabling the visualization of many of the protepglycans within the ECM of decellularized tissues [55]. Cartilage ECM is composed of type II collagen, elastic fibers, proteoglycans, hyaluronic acid, and other macromolecules [56]. The ECM plays a critical role in directing cellular behaviors and resisting the forces generated as these cartilage elements differentiate into bones, with a small portion of cartilage remaining on the articulating surfaces. When these complex tissues are damaged, in particular the articular cartilage, the functionality is difficult to recover. Consequently, there is a significant focus on generating scaffold materials that can restore the function and structure of adult skeletal tissues by recapitulating the environment found during development. However, the structure of cartilage matrix remains elusive as the majority of the ECM networks are unresolvable using conventional diffraction-limited microscopy. Here we used INSPR to reconstruct a 3D super-resolution volume of a 20-μm-thick developing cartilage tissue with an axial depth of 14 μm. Despite the imaging depth and complex tissue environment, INSPR resolved fine elastin-based, elastic fibers in three dimensions among the proteoglycans (Figures 6A and 6B, Movie S7). These elastic fibers, independent of their orientations, were resolved (Figures 6C–6F) with a lateral width quantified by FWHM from 58 nm to 194 nm (Figure 6G, 109±33 nm, 60 measurements, profile positions shown in Figure S5E) and an axial width from 78 to 281 nm (Figure 6H, 160±55 nm, 40 measurements). We observed that these elastic fibers evolved along their paths in the tissue with the diameter changing as much as 80% (defined by FWHM_max_/ FWHM_min_ – 1, average of 41%, 15 measurements, Figure 6G) in the lateral plane, an observation in agreement with previous studies using electron microscopy in adult articular cartilage and skin [57]. In addition, the reconstructed super-resolution volume using INSPR allows us to trace individual elastic fibers in three dimensions within the tissue while observing their dynamic size changes along the path. The observation of nanoscale architectures of specific extracellular matrix constituents will help in designing suitable regenerative scaffolds to restore the functionality to cartilage and other damaged tissues.

**Figure 6.**
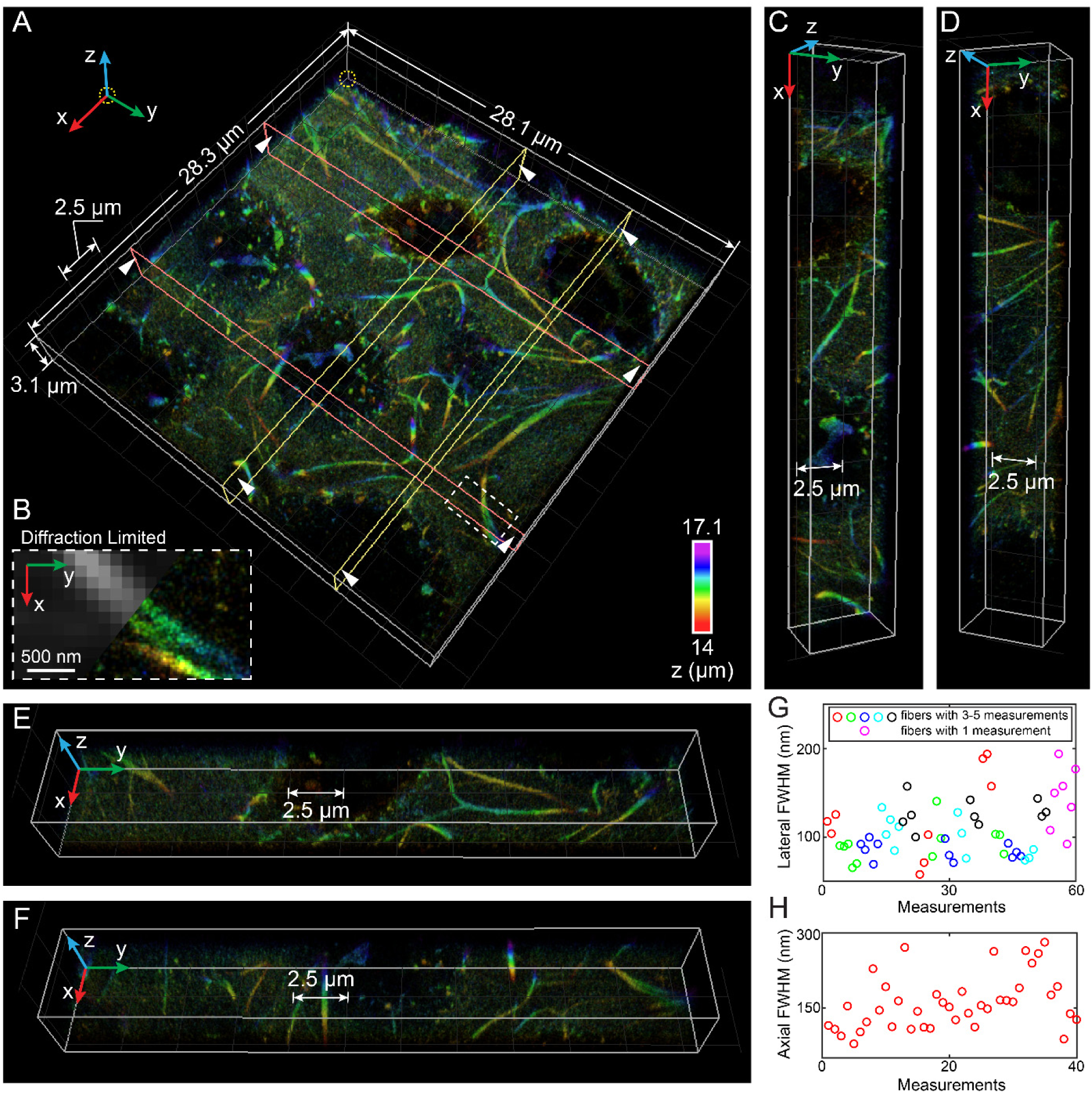
3D super-resolution reconstruction of ECM in developing cartilage within mouse forelimbs using INSPR. (A) 3D overview of a 3.1-µm-thick volume reconstruction of cartilage tissue at a depth of 14 µm from the coverslip surface, where fine elastic fibers are resolved with distinctly traceable orientations in three dimensions among the proteoglycans. An animated 3D reconstruction is shown in Movie S7. (B) Enlarged x-y view of the area as indicated by the white dashed boxed region in (A), showing the details of a split elastic fiber (right), which is not resolvable when using conventional diffraction-limited microscopy (left). (C–F) Cross sections of the reconstruction in (A). (G) Distribution of lateral FWHM, measured from 15 long fibers (3–5 measurements per fiber, indicated by red, green, blue, cyan, and black circles, where adjacent circles with the same color refer to multiple measurements in one fiber) and 7 short fibrils (single measurement per fiber, indicated by magenta circles), both in the x-y plane. Lateral FWHM varies from 58 to 194 nm, with a mean of 109 nm and a standard deviation of 33 nm. (H) Distribution of axial FWHM, measured from 40 typical elastic fibers in the x-z plane. Axial FWHM varies from 78 to 281 nm, with a mean of 160 nm and a standard deviation of 55 nm.

## DISCUSSION

We demonstrated INSPR in retrieving accurate *in situ* 3D PSF responses of single fluorescent emitters directly from single molecule datasets, a significant advancement from previous *in vitro* methods. INSPR enables us to image whole cells and tissues with 20–30 nm lateral and 40–80 nm axial resolution throughout their entire volume with high reconstruction accuracy. We demonstrated this capacity through resolving nanoscale details of mitochondrial networks, microtubules (Figure S6), and nuclear pore complexes of whole cells in 2D and 3D cultures, amyloid β plaques and fibrils in brain slices, and developing cartilage ECM in mouse forelimbs.

Our experiments in cell and tissue specimens demonstrate the capability of INSPR to image at a depth of <20 μm. However, imaging above this demonstrated depth will be challenged by the constantly decreasing information content (Fisher information, **Supplementary Notes**) of single molecule emission patterns due to aberrations. Such information loss cannot be recovered by algorithm but rather requires a physical element that modifies the distorted wavefront prior to the detection of the fluorescent signal. The combination of adaptive optics (AO) [29,30,58–64] with INSPR will allow restoring emission pattern information and pin-pointing 3D location of single molecules with high accuracy simultaneously. In addition, INSPR can be combined with light-sheet illumination approaches [65–67] and tissue clearing and expansion methods [68–70] to further reduce the fluorescence background and increase the achievable resolution, therefore opening doors to observe the static and dynamic conformation of complex intra- and extra-cellular constituents over large tissue volumes at the nanoscale level.

Nanoscopy of specimens that are living, of high throughput, and with multi-color probes will induce time-, region-, and channel-dependent aberrations in single molecule datasets. Future applications of INSPR will also allow extraction of such temporally, spatially, and spectrally varying single molecule 3D responses to ensure pin-pointing single molecule positions with high precision and accuracy. Therefore, we expect INSPR will be a useful tool in visualizing cellular structures and protein functions throughout whole cells and tissues across diverse biological and biomedical model systems.

## AUTHOR CONTRIBUTIONS

F.X., D.M., and F.H. conceived the project. F.X., S.L., and F.H. developed the algorithm. F.X. wrote the software and performed simulation. D.M. developed the microscope setup and performed super-resolution experiments. F.X., D.M., and S.L. visualized experimental data. D.M., K.P.M., Y.B., Y.W., C.B., T.K., P.Y., S.C., G.E.L., and F.H. designed the experiments. D.M., K.P.M., Y.B., Y.W., C.B., and T.K. prepared biological samples. P.Y., S.C., G.E.L., and F.H. supervised the study. All authors wrote the manuscript.

## ACKNOWLEGEMENTS

We would like to thank Karthigeyan Dhanasekaran and Patrick Lusk (Yale University) for sharing the labeling protocol of Nup98 and interpretation of the resolved Nup98 structures. We thank Michael J. Mlodzianoski for initial instrument design, Sha An for help in instrument alignment, and David A. Miller for providing labeling protocols of mitochondria and microtubules. F.X., D.M., S.L., C.B., and F.H. were supported by grants from NIH (GM119785) and DARPA (D16AP00093). K.P.M. and G.E.L were supported by grant from NIH (AG051495 and AG050597). Y.B. and S.C. were supported by grant from NIH (AR071359). Y.W. and P.Y. were supported by grant from NIH (1R01EB018659) and Harvard Medical School Dean’s Initiative Grant. Data that support the findings of this study are available from the corresponding author upon request.

