## Supplementary Materials for "Three dimensional nanoscopy of whole cells and tissues with *in situ* point spread function retrieval"

**This PDF file includes:**

Supplementary Figures S1 to S6

Supplementary Tables S1 to S2

Supplementary Notes

Additional References

**Other supplementary materials for this manuscript include the following:**

**Movie S1**: Performance demonstration of INSPR on simulated single molecule emission patterns.

**Movie S2**: INSPR estimates wavefront distortions induced by deformable mirror from single molecule dataset of immunolabeled TOM20.

**Movie S3**: 3D super-resolution reconstruction of mitochondrial networks in fixed COS-7 cells.

**Movie S4**: 3D super-resolution reconstruction of nuclear pore complex protein Nup98 on an entire cell nucleus in fixed COS-7 cells.

**Movie S5**: 3D super-resolution reconstruction of amyloid β plaque with low fibril density in a 30-μm-thick brain section of a mouse model of Alzheimer's disease.

**Movie S6**: 3D super-resolution reconstruction of amyloid β plaque with high fibril density in a 30-μm-thick brain section of a mouse model of Alzheimer's disease.

**Movie S7**: 3D super-resolution reconstruction of elastic fibers of developing cartilage in embryonic mouse forelimbs.

### Supplementary Figures

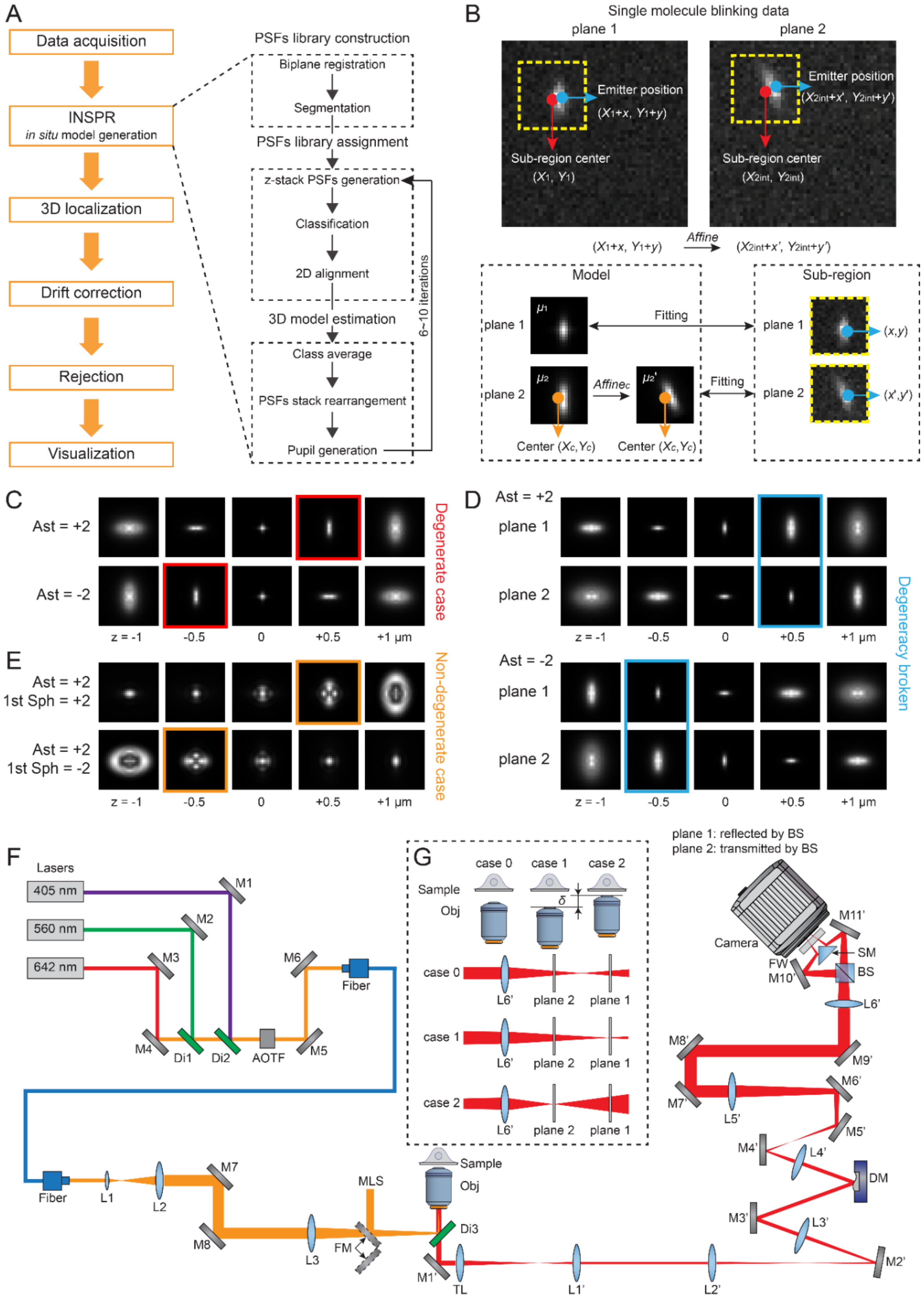

**Figure S1.** INSPR framework, degeneracy demonstration, and setup diagram.

(A) INSPR framework and detailed process of three key components: PSFs library construction, PSFs library assignment, and 3D model estimation.

(B) Channel-specific localization. Instead of transforming data directly from plane 2 to plane 1, we transform the identified sub-region center from plane 1 $(X_1, Y_1)$ to plane 2 $(X_{2\text{int}}, Y_{2\text{int}})$, and crop two sub-regions from both planes. We first generate a fitting model $\mu_1$ for plane 1, and then generate a channel-specific model $\mu_2{}'$ for plane 2 which shares the same shape information with the cropped sub-region in plane 2. After that, this pair of channel-specific models is used for single molecule localization.

(C) Degeneracy exists in the single plane configuration, where positive (top row) and negative (bottom row) vertical astigmatism aberrations generate identical PSFs at opposite axial positions (as highlighted by the red-boxed regions). Ast: vertical astigmatism.

(D) Degeneracy is broken in the biplane configuration, where positive (top row) and negative (bottom row) vertical astigmatism aberrations generate different PSF pairs at opposite axial positions (as highlighted by the blue-boxed regions).

(E) Degeneracy is absent in the single plane configuration when using a cylindrical lens with prior knowledge such as positive vertical astigmatism, where positive (top row) and negative (bottom row) primary spherical aberrations generate different PSFs at opposite axial positions (as highlighted by the orange-boxed regions). 1st Sph: primary spherical.

(F) Diagram of the microscope setup. M1–M8: mirrors in the excitation path; Di1–Di3: dichroic mirrors; AOTF: acousto-optic tunable filter; L1–L3: lenses in the excitation path; FM: motorized flip mirror; MLS: mercury light source; Obj: objective lens; M1'–M11': mirrors in the emission path; TL: tube lens; L1'–L6': lenses in the emission path; DM: deformable mirror; BS: 50/50 non-polarizing beam splitter cube; SM: 90° specialty mirror; FW: motorized filter wheel. Focal lengths of lenses are, L1: 20 mm, L2: 125 mm, L3: 400 mm, Obj: 1.8 mm, TL: 180 mm, L1': 88.9 mm, L2': 250 mm, L3': 400 mm, L4': 150 mm, L5': 500 mm, L6': 250 mm.

(G) Definition of biplane distance. In case 0, the focal plane is in the middle of plane 1 and plane 2. In case 1, the objective lens is moved down to make plane 1 in focus. In case 2, the objective lens is moved up to make plane 2 in focus. The axial movement of the objective lens from case 1 to case 2 is defined as biplane distance δ.

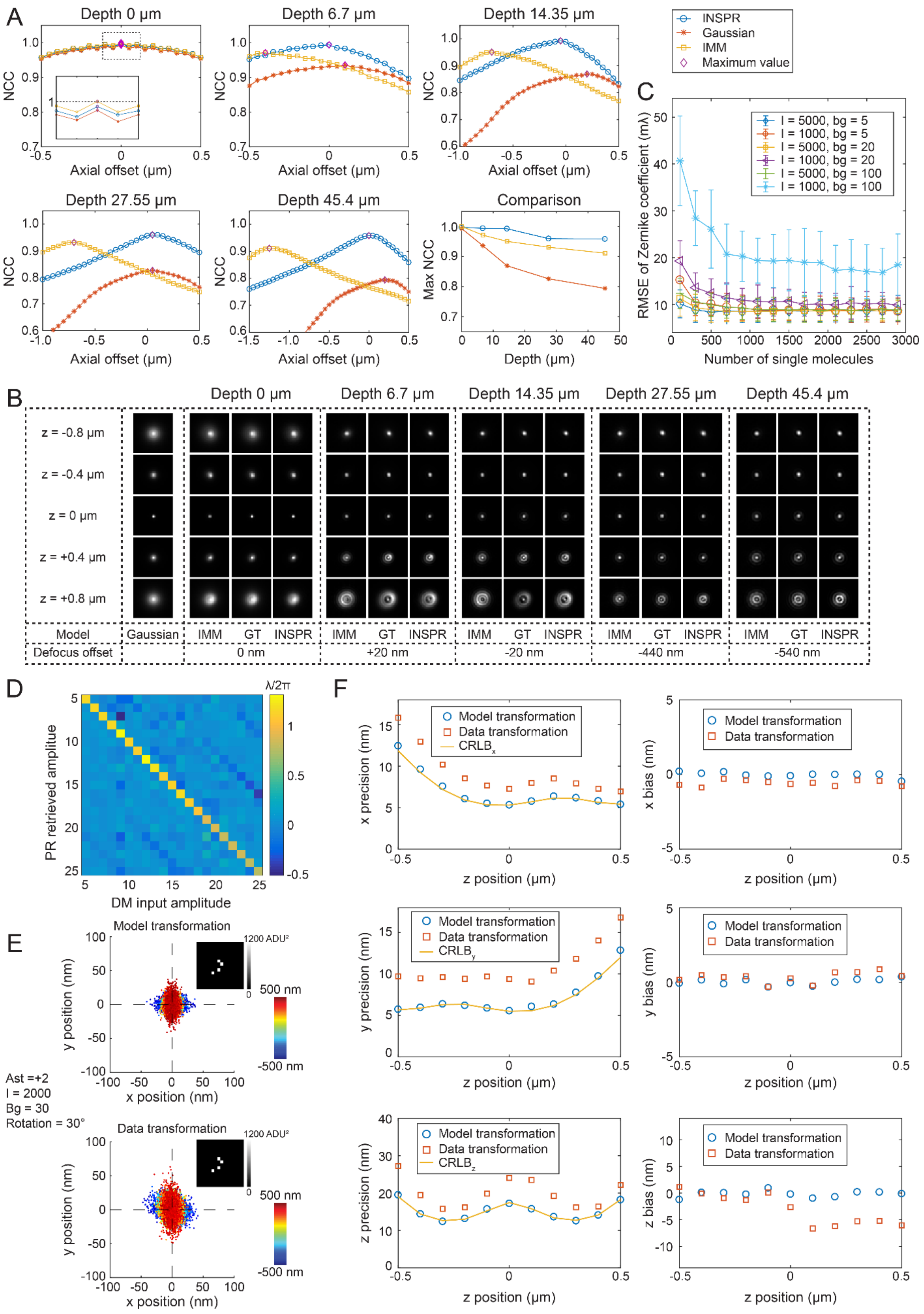
A
Depth 0 μm
Depth 6.7 μm
Depth 14.35 μm
Depth 27.55 μm
Depth 45.4 μm
Comparison
NCC
Max NCC
Axial offset (μm)
Depth (μm)
INSPR
Gaussian
IMM
Maximum value
C
RMSE of Zernike coefficient (mλ)
Number of single molecules
I = 5000, bg = 5
I = 1000, bg = 5
I = 5000, bg = 20
I = 1000, bg = 20
I = 5000, bg = 100
I = 1000, bg = 100
B
z = -0.8 μm
z = -0.4 μm
z = 0 μm
z = +0.4 μm
z = +0.8 μm
Model
Gaussian
IMM GT INSPR
Defocus offset
0 nm
+20 nm
-20 nm
-440 nm
-540 nm
D
λ/2π
PR retrieved amplitue
DM input amplitude
E
Model transformation
Data transformation
1200 ADU²
500 nm
-500 nm
y position (nm)
x position (nm)
Ast =+2
I = 2000
Bg = 30
Rotation = 30°
F
Model transformation
Data transformation
CRLB$_x$
CRLB$_y$
CRLB$_z$
x precision (nm)
y precision (nm)
z precision (nm)
x bias (nm)
y bias (nm)
z bias (nm)
z position (μm)

**Figure S2.** Performance quantification of *in situ* PSF retrieval with INSPR.

(A) Similarity of the 3D PSFs at different imaging depths when using INSPR (blue circles), Gaussian model (orange stars), and theoretical index mismatch model (IMM, yellow squares). For each depth, the reference 3D PSFs were simulated according to our previously measured aberrations with an axial range from –800 to +800 nm and a step size of 100 nm. Three models were then used to generate 3D PSFs in the same axial range with different axial offsets. For each depth, 3D normalized cross correlation (NCC) coefficients between the reference PSFs and PSFs generated by three models at different axial offsets are shown with the maximum values marked (purple diamonds). At the depth of 0 μm, the maximum NCC of INSPR is closer to that of IMM, compared with Gaussian. As the depth increases, it remains around 0.95, higher than those of Gaussian and IMM.

(B) 3D PSFs when using different models such as Gaussian, INSPR, and IMM in comparison to ground truth (GT). PSFs shown are chosen from the maximum NCC (purple diamonds) for each depth in (A). The PSFs retrieved by INSPR show higher similarity with ground truth, in comparison to those retrieved by IMM. The defocus offset (i.e., the axial shift from the actual focal plane) was obtained by finding the maximum-intensity plane of ground truth PSFs along the axial direction.

(C) Root-mean-square error (RMSE) of Zernike coefficients between the *in situ* model and ground truth in different photon and background conditions. In each condition, data were simulated with Zernike coefficients randomly sampled from –1 to +1 (unit: λ/2π) in each trial (11 trials in total). As the number of single molecules increases, RMSE converges to the stable estimation. High signal to background ratio (SBR) cases require less than 300 single molecules, while low SBR cases require more than 2100 single molecules.

(D) Heat map showing the relationship between the input and phase retrieved (PR) amplitudes of 21 Zernike modes. The largest estimated amplitude is along the diagonal elements in the response matrix. The error between the input and output amplitudes along the diagonal elements is 6% in the first 14 Zernike modes and 27% in the last 7 Zernike modes (details in 'Calibration of deformable mirror' section).

(E) Scatter plots of lateral localizations using model transformation (top) and data transformation (bottom) for PSFs with vertical astigmatism. Method using model transformation provides localization precision achieving the CRLB and smaller bias. For each PSF, the total photon counts $I$ was set to 2000, and the background $bg$ was set to 30. Plane 1 and plane 2 are related with an affine transformation including a rotation of 30 degrees. Both Poisson noise and the pixel-dependent sCMOS readout noise (the variance distribution is shown in the inset) were considered in simulated datasets.

(F) Localization precisions and biases in the x, y, and z dimensions for datasets in (E). The precisions of model transformation are close to CRLB in all three dimensions, and its biases are smaller than those of data transformation.

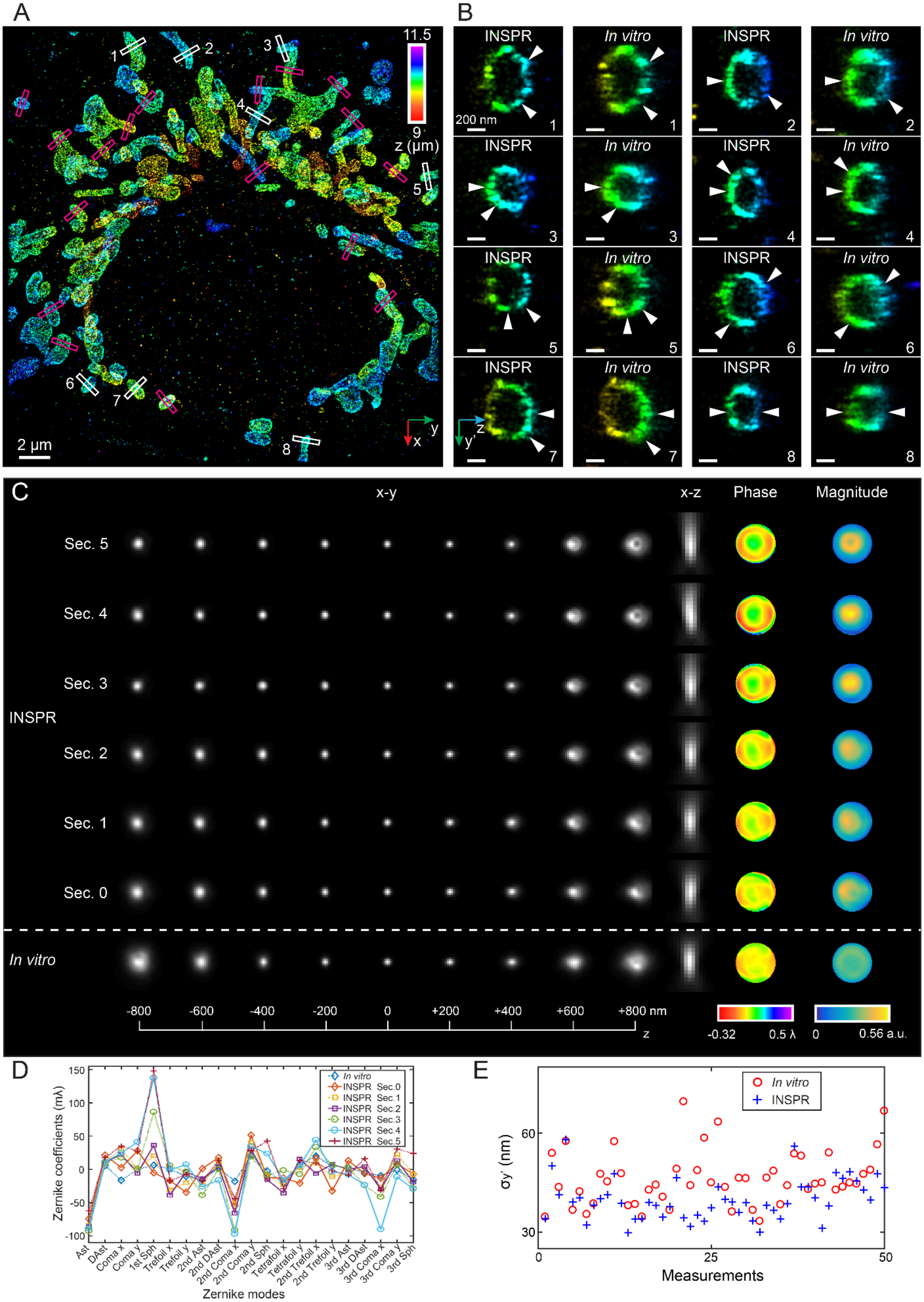

A
B
C
D
E
INSPR
In vitro
x-y
x-z
Phase
Magnitude
Sec. 5
Sec. 4
Sec. 3
Sec. 2
Sec. 1
Sec. 0
2 μm
200 nm
z (μm)
11.5
9
-800
-600
-400
-200
0
+200
+400
+600
+800 nm
-0.32
0.5 λ
0
0.56 a.u.
Zernike coefficients (mλ)
Zernike modes
σy' (nm)
Measurements

**Figure S3.** 3D reconstruction of TOM20 in COS-7 cells using INSPR.

(A) x-y overview of the mitochondrial network showing the positions of 25 typical outer membrane contours as indicated by the magenta and white boxed regions.

(B) Enlarged y'-z views of the outer membrane structures as indicated by the white-boxed regions in (A), showing the reconstructed images using INSPR (left) and the *in vitro* (right) approach. Here the orientation of the cross section is rotated to allow projection of the 3D membrane bounded structures to the 2D image.

(C) x-y and x-z views of the PSFs generated by the INSPR models of different sections (top rows) and those generated by the *in vitro* model (bottom row), as well as the phase and magnitude of the corresponding pupils. As the imaging depth increases, the axially stretched PSFs retrieved by INSPR show the existence of sample-induced aberrations, which cannot be characterized by the *in vitro* model. Sec.: section.

(D) Amplitudes of 21 Zernike modes (Wyant order, from vertical astigmatism to tertiary spherical aberration) decomposed from the INSPR and *in vitro* pupils. As the imaging depth increases, the INSPR pupils and their corresponding Zernike amplitudes show a progressive increase of major aberrations such as spherical and coma, while the PSF retrieved from fluorescent beads can only characterize instrument imperfections but fails to reflect sample-induced aberrations and their depth-dependent variations due to its *in vitro* nature.

(E) Distribution of $\sigma_{y'}$ obtained from the intensity profiles of 25 typical outer membranes in (A), which demonstrates an obvious improvement in the lateral resolution achieved by INSPR (blue plus signs) in comparison to the *in vitro* approach (red circles).

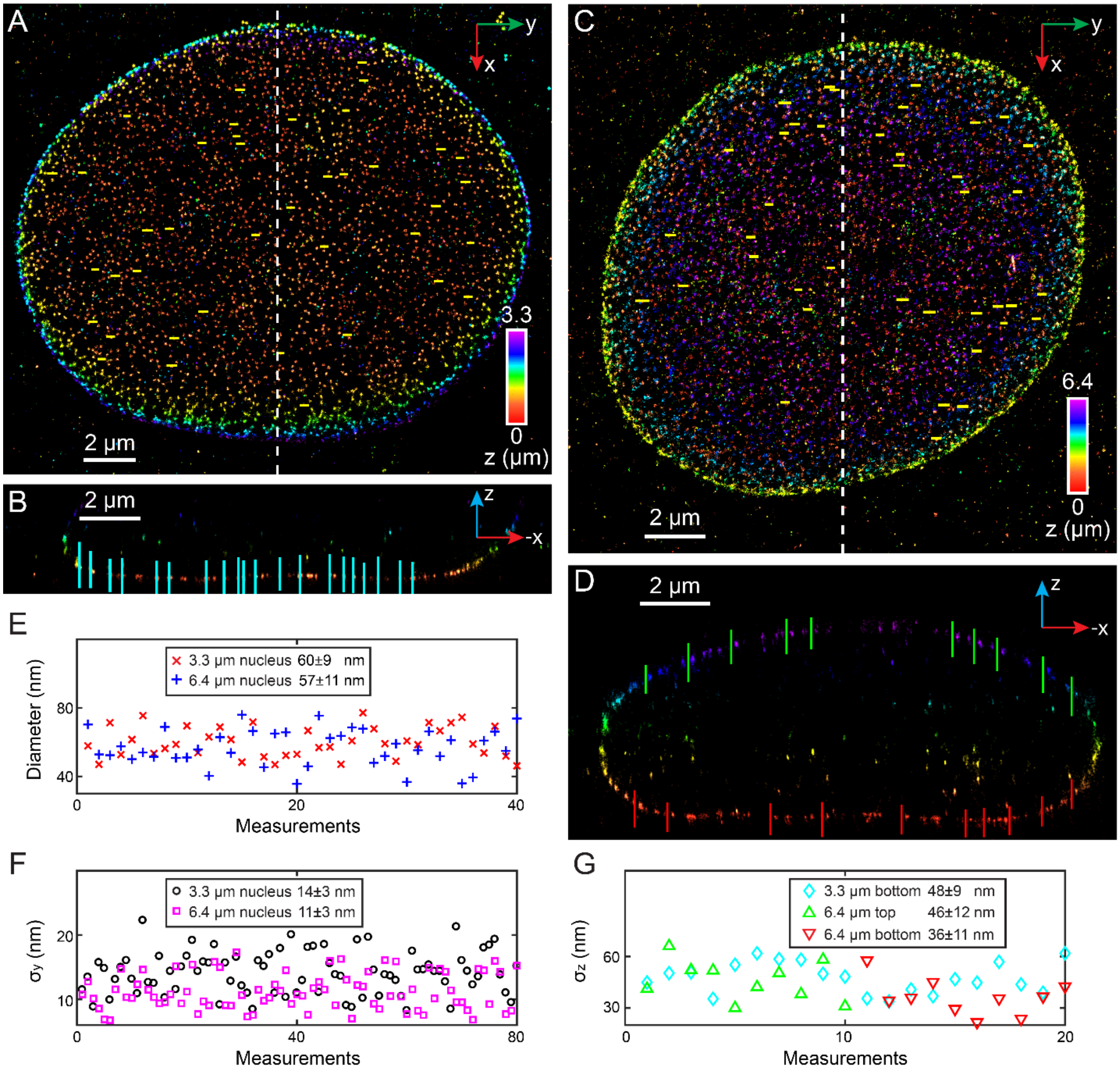

**Figure S4.** 3D reconstruction of nuclear pore complex protein Nup98 in COS-7 cells using INSPR.

(A) x-y overview of the 3.3-μm-thick section of the nucleus showing the positions of 40 typical resolved Nup98 structures as indicated by the yellow lines.

(B) x-z slice along the white dashed line in (A), showing the positions of 20 typical Nup98 structures as indicated by the cyan lines. The thickness along the y direction is 500 nm.

(C) x-y overview of the 6.4-μm-thick entire nuclear envelope showing the positions of 40 typical resolved Nup98 structures as indicated by the yellow lines.

(D) x-z slice along the white dashed line in (C), showing the positions of 10 Nup98 structures on the top surface (green lines) and 10 typical Nup98 structures on the bottom surface (red lines). The thickness along the y direction is 500 nm.

(E) Distribution of diameters measured from 40 resolved Nup98 structures in the x-y plane shown in (A) and (C). The average diameter is 60±9 nm for the 3.3-μm-thick nucleus (red crosses), and 57±11 nm for the 6.4-μm-thick nucleus (blue plus signs).

(F) Distribution of $\sigma_y$ measured from 40 resolved Nup98 structures in the x-y plane shown in (A) and (C). The average $\sigma_y$ is 14±3 nm for the 3.3-μm-thick nucleus (black circles), and 11±3 nm for the 6.4-μm-thick nucleus (magenta squares).

(G) Distribution of $\sigma_z$. For the 3.3-μm-thick nucleus, the average $\sigma_z$ is 48±9 nm (cyan diamonds), measured from 20 Nup98 structures in the x-z plane shown in (B). For the 6.4-μm-thick nucleus, the average $\sigma_z$ is 46±12 nm for the top surface (green upward-pointing triangles), measured from 10 Nup98 structures as indicated by the green lines in (D); and 36±11 nm for the bottom surface (red downward-pointing triangles), measured from 10 Nup98 structures as indicated by the red lines in (D).

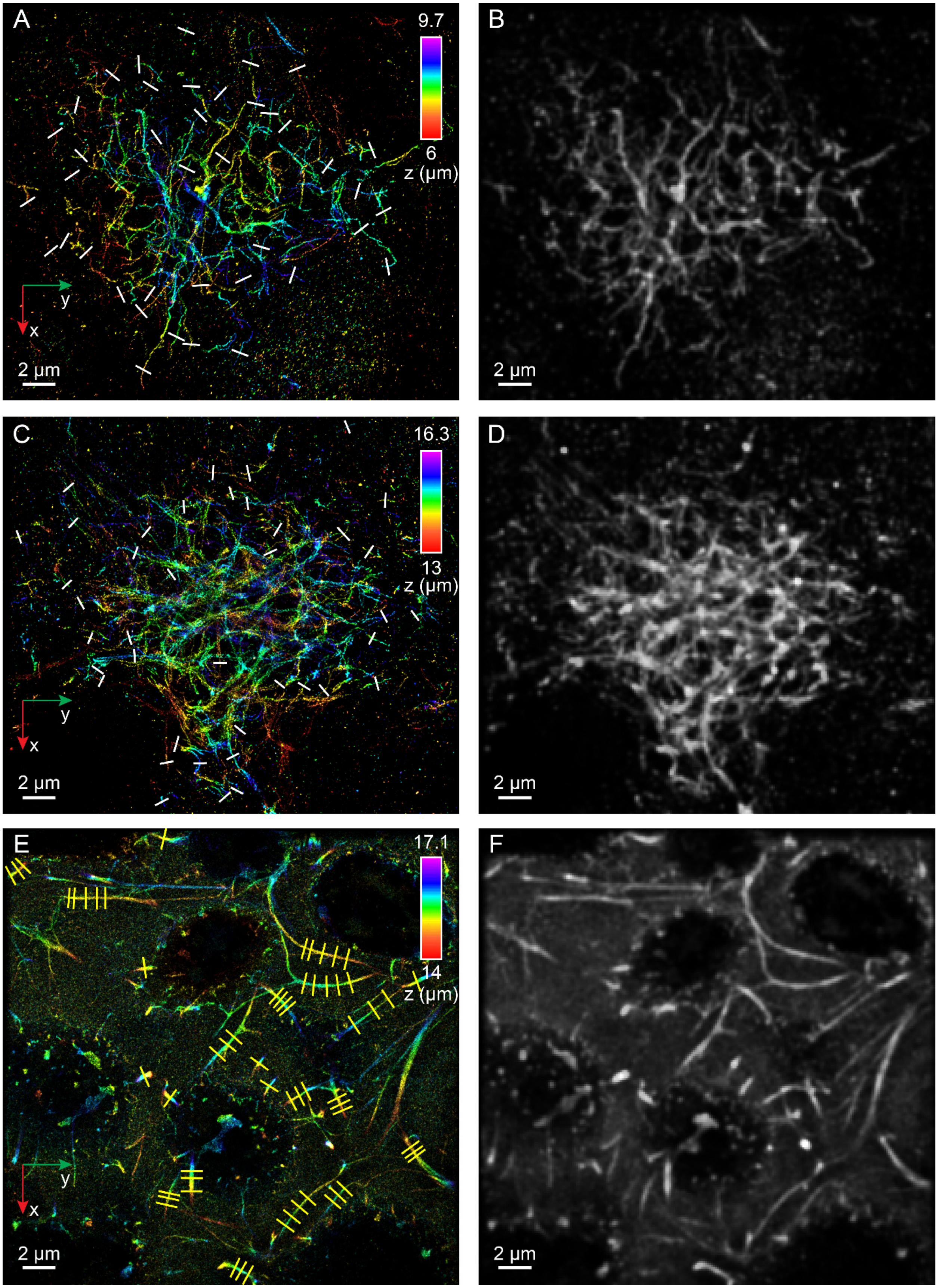
A
9.7
6
z (μm)
y
x
2 μm
B
2 μm
C
16.3
13
z (μm)
y
x
2 μm
D
2 μm
E
17.1
14
z (μm)
y
x
2 μm
F
2 μm

**Figure S5.** 3D reconstruction of amyloid β fibrils in mouse brain sections and developing cartilage in mouse forelimbs using INSPR.

(A) x-y overview of the super-resolution volume of a low-density plaque with a relatively low density of amyloid β fibrils showing the positions of 40 typical individual fibrils as indicated by the white lines.

(B) Diffraction-limited image of (A).

(C) x-y overview of the super-resolution volume of a high-density plaque with a relatively high density of amyloid β fibrils showing the positions of 40 typical individual fibrils as indicated by the white lines.

(D) Diffraction-limited image of (C).

(E) x-y overview of the 3.1-µm-thick developing cartilage showing the positions of 15 long elastic fibers with 3–5 measurements per fiber and 7 short elastic fibers with single measurement, as indicated by the yellow lines.

(F) Diffraction-limited image of (E).

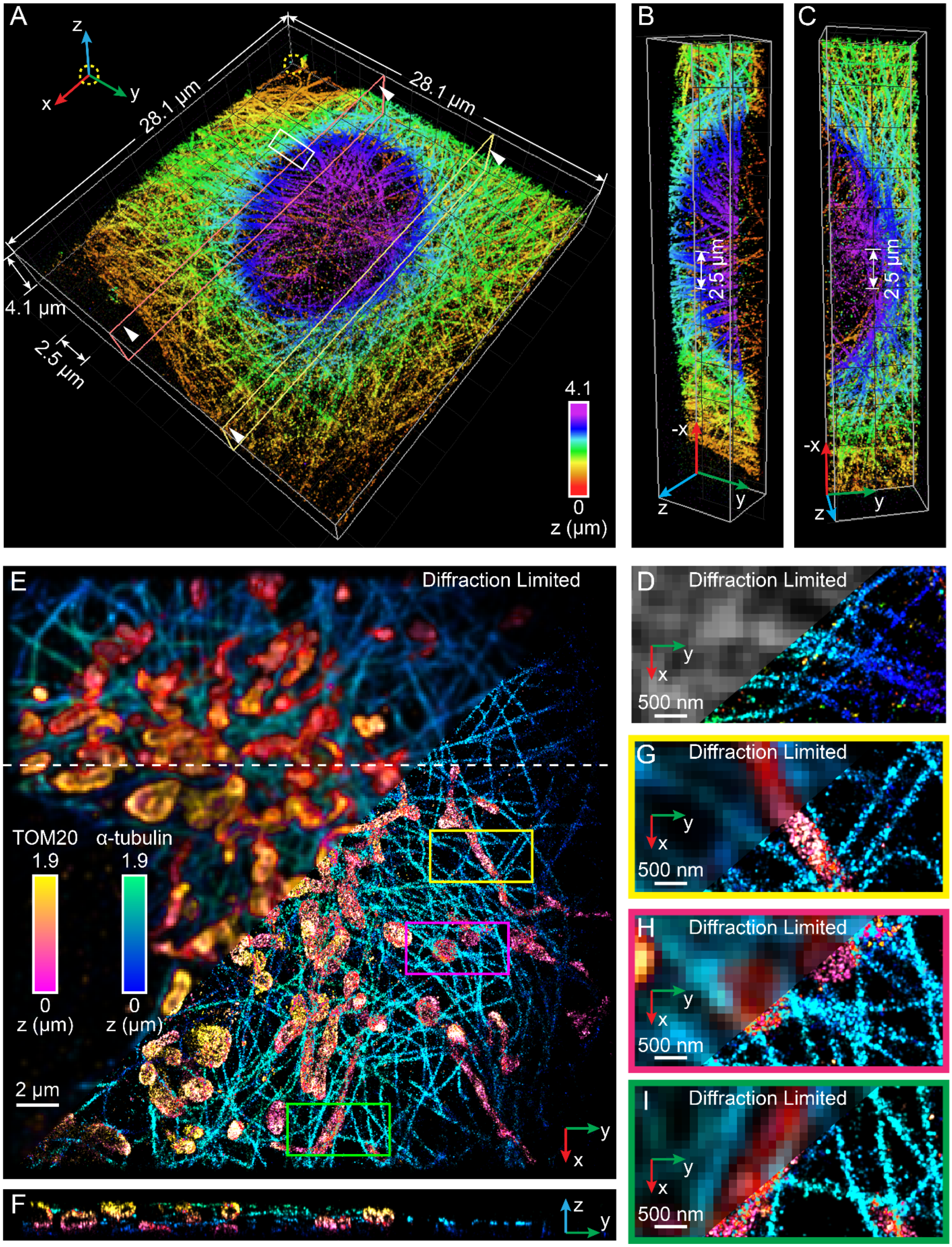
A
z
x
y
28.1 µm
28.1 µm
4.1 µm
2.5 µm
4.1
0
z (µm)
B
2.5 µm
-x
z
y
C
2.5 µm
-x
z
y
E
Diffraction Limited
TOM20
α-tubulin
1.9
1.9
0
0
z (µm)
z (µm)
2 µm
y
x
F
z
y
D
Diffraction Limited
y
x
500 nm
G
Diffraction Limited
y
x
500 nm
H
Diffraction Limited
y
x
500 nm
I
Diffraction Limited
y
x
500 nm

**Figure S6.** INSPR reconstruction of microtubules in collagen embedded 3D-cultured cells and TOM20 co-stained with α-tubulin using Exchange-PAINT in 2D cultures.

(A) 3D super-resolution image overview of microtubules in collagen embedded 3D cultured cells with a total thickness of 4.1 μm.

(B and C) Cross sections of the super-resolution reconstruction as indicated by the orange (B) and yellow (C) planes in (A).

(D) Enlarged view of the area as indicated by the white-boxed region in (A), showing fine and dense microtubules (right) and its counterpart under the diffraction limit (left).

(E and F) Reconstruction of mitochondria (TOM20) and microtubules (α-tubulin) in a COS-7 cell immuno-labeled with DNA-PAINT. An x-y overview (E) and y-z slice along the white dashed line (F) are shown. The thickness of the axial slice along the x direction is 200 nm.

(G–I) Enlarged views of the areas as indicated by the yellow (G), magenta (H), and green (I) boxed regions in (E), showing interactions between microtubules and mitochondria (right) and their counterparts under the diffraction limit (left).

### Supplementary Tables

| Figures and Panels | Frames per step | Number of steps | Number of cycles | Total frames | ROI thickness (µm) | FOV center depth (µm) | Mean lateral localization precision (nm) | Mean axial localization precision (nm) | Mean photon counts | Mean background count | Number of localizations |
|---|---|---|---|---|---|---|---|---|---|---|---|
| Figure 3 TOM20 | 2000 | 6 | 20 | 240,000 | ~2.5 | ~10.3 | 8.2 (x) 7.2 (y) | 21.3 | 5951 | 144.2 | 1,052,610 |
| Figure 4A Nup98 | 2000 | 5 | 10 | 100,000 | ~3.3 | ~1.7 | 9.2 (x) 10.4 (y) | 33.8 | 2531 | 34.2 | 622,142 |
| Figure 4E Nup98 | 1000 | 14 | 10 | 140,000 | ~6.4 | ~3.2 | 9.3 (x) 10.4 (y) | 35.0 | 2498 | 34.9 | 381,054 |
| Figure 5A Amyloid β | 2000 | 5 | 9 | 90,000 | ~3.7 | ~7.9 | 7.2 (x) 7.3 (y) | 28.9 | 5419 | 99.2 | 898,404 |
| Figure 5K Amyloid β | 2000 | 5 | 14 | 140,000 | ~3.3 | ~14.7 | 8.7 (x) 8.6 (y) | 32.8 | 4138 | 104.2 | 1,090,760 |
| Figure 6 Cartilage | 2000 | 5 | 13 | 130,000 | ~3.1 | ~15.6 | 11.4 (x) 11.1 (y) | 44.6 | 3043 | 138.5 | 1,988,098 |
| Figure S6A α-tubulin | 2000 | 7 | 10 | 140,000 | ~4.1 | ~2.1 | 7.2 (x) 7.4 (y) | 28.0 | 2849 | 39.1 | 1,459,461 |
| Figure S6E α-tubulin | 2000 | 1 | 46 | 92,000 | ~1.7 | ~0.9 | 8.0 (x) 8.6 (y) | 35.0 | 2497 | 38.8 | 1,876,043 |
| Figure S6E TOM20 | 2000 | 1 | 50 | 100,000 | ~1.9 | ~1.0 | 7.8 (x) 7.9 (y) | 36.5 | 2956 | 54.9 | 1,410,818 |

**Table S1**. Imaging parameters for experimental data.

All images were recorded with a frame rate of 50 Hz and a piezo step size of 400 nm for multi-section imaging. Related to Figures 3–6 and Figure S6. ROI: region of interest; FOV: field of view. Lateral and axial localization precisions were estimated by Cramér-Rao lower bound (CRLB).

| Zernike-based aberrations | Amplitude (λ/2π) | $I_{\mathrm{seg}}$ (photons) | Number of sub-regions | $\mathrm{Sim}_{\mathrm{min}}$ | $N_g$ | Z_shift_mode |
|---|---|---|---|---|---|---|
| Ast | +1 | 80 | 5025 | 0.5 | 25 | Shift |
| | –1 | 80 | 4978 | 0.5 | 25 | Shift |
| DAst | +1 | 80 | 4948 | 0.5 | 25 | Shift |
| | –1 | 85 | 4933 | 0.5 | 25 | Shift |
| Coma x | +1 | 70 | 5053 | 0.5 | 25 | Shift |
| | –1 | 85 | 4990 | 0.5 | 25 | Shift |
| Coma y | +1 | 75 | 4959 | 0.5 | 25 | Shift |
| | –1 | 70 | 4888 | 0.5 | 25 | Shift |
| 1st Sph | +1 | 45 | 4831 | 0.6 | 25 | No shift |
| | –1 | 35 | 4829 | 0.6 | 25 | No shift |
| Trefoil x | +1 | 50 | 5114 | 0.5 | 25 | Shift |
| | –1 | 50 | 5069 | 0.5 | 25 | Shift |
| Trefoil y | +1 | 45 | 5088 | 0.5 | 25 | Shift |
| | –1 | 40 | 5160 | 0.5 | 25 | Shift |
| 2nd Ast | +1 | 35 | 5058 | 0.6 | 15 | Shift |
| | –1 | 45 | 4959 | 0.6 | 15 | Shift |
| 2nd DAst | +1 | 45 | 5145 | 0.6 | 25 | Shift |
| | –1 | 50 | 5110 | 0.6 | 25 | Shift |
| 2nd Coma x | +1 | 60 | 5055 | 0.5 | 15 | No shift |
| | –1 | 55 | 4942 | 0.5 | 15 | No shift |
| 2nd Coma y | +1 | 45 | 5157 | 0.5 | 25 | No shift |
| | –1 | 60 | 5155 | 0.5 | 25 | No shift |
| 2nd Sph | +1 | 60 | 4879 | 0.6 | 25 | No shift |
| | –1 | 90 | 5067 | 0.5 | 15 | No shift |
| Tetrafoil x | +1 | 65 | 5020 | 0.6 | 25 | No shift |
| | –1 | 60 | 5001 | 0.6 | 25 | No shift |
| Tetrafoil y | +1 | 60 | 5032 | 0.5 | 25 | No shift |
| | –1 | 55 | 5036 | 0.5 | 25 | No shift |
| 2nd Trefoil x | +1 | 35 | 5053 | 0.5 | 25 | No shift |
| | –1 | 40 | 4815 | 0.5 | 25 | No shift |
| 2nd Trefoil y | +1 | 35 | 4770 | 0.5 | 25 | No shift |
| | –1 | 35 | 5073 | 0.5 | 25 | No shift |
| 3rd Ast | +1 | 40 | 5045 | 0.5 | 25 | No shift |
| | –1 | 40 | 4935 | 0.5 | 25 | No shift |
| 3rd DAst | +1 | 45 | 5126 | 0.5 | 25 | No shift |
| | –1 | 45 | 4839 | 0.5 | 25 | No shift |
| 3rd Coma x | +1 | 60 | 4880 | 0.5 | 25 | No shift |
| | –1 | 50 | 4949 | 0.5 | 25 | No shift |
| 3rd Coma y | +1 | 85 | 5021 | 0.5 | 25 | No shift |
| | –1 | 80 | 4941 | 0.5 | 25 | No shift |
| 3rd Sph | +1 | 75 | 5083 | 0.6 | 25 | No shift |
| | –1 | 65 | 5115 | 0.6 | 25 | No shift |

**Table S2.** Parameter settings of INSPR for performance test based on wavefront distortions induced by a deformable mirror.

Common parameters used for INSPR include the sub-region size of 40 × 40 pixels, $d_{\mathrm{thresh}}$ of 28, $I_{\mathrm{init}}$ of 30, $d_t$ of 145 nm, and XY_shift_mode of 'together shift'.

Ast: vertical astigmatism; DAst: diagonal astigmatism; Coma x: horizontal coma; Coma y: vertical coma; 1st Sph: primary spherical; Trefoil x: horizontal trefoil; Trefoil y: vertical trefoil; 2nd Ast: secondary vertical astigmatism; 2nd DAst: secondary diagonal astigmatism; 2nd Coma x: secondary horizontal coma; 2nd Coma y: secondary vertical coma; 2nd Sph: secondary spherical; Tetrafoil x: horizontal tetrafoil; Tetrafoil y: vertical tetrafoil; 2nd Trefoil x: secondary horizontal trefoil; 2nd Trefoil y: secondary vertical trefoil; 3rd Ast: tertiary horizontal astigmatism; 3rd DAst: tertiary diagonal astigmatism; 3rd Coma x: tertiary horizontal coma; 3rd Coma y: tertiary vertical coma; 3rd Sph: tertiary spherical.

### Supplementary Notes

#### Preparation of fluorescent beads on coverslips

25-mm-diameter coverslips (Bioscience Tools, Cat#CSHP-No1.5-25) were cleaned successively in ethanol (Decon, Cat#2701) and HPLC grade water (Fisher Chemical, Cat#W5-4) three times, and then dried with compressed air. 100-nm-diameter crimson beads (Invitrogen, custom-designed) were diluted to 1:100,000 in deionized water. 200 μL of poly-L-lysine solution (Sigma-Aldrich, Cat#P4707) was added on the coverslip, incubated for 20 min and subsequently rinsed with deionized water. 200 μL of diluted bead solution was added on the center of the coverslip and was incubated for 20 min at room temperature (RT). The coverslip was subsequently rinsed with deionized water and drained. 20 μL of 38% TDE (Sigma-Aldrich, Cat#166782) in PBS (Gibco, Cat#10010023) was added on the center of the coverslip. Another 25-mm-diameter coverslip (also cleaned by using the above protocol) was placed on top of this coverslip. This coverslip sandwich was sealed with melted valap (1:1:1 [w/w/w] mixture of lanolin, paraffin, and Vaseline (Sigma-Aldrich, Cat#L7387, 18634, and 16415)).

#### Cell culture

COS-7 cells (ATCC, Cat#CRL-1651) were immuno-labeled with TOM20, α-tubulin, and Nup98. COS-7 cells were grown on coverslips in 6-well plates and cultured in DMEM (ATCC, Cat#30-2002) with 10% FBS (ATCC, Cat#30-2020) and 1% Penicillin-Streptomycin (Gibco, Cat#15140122) at 37 °C with 5% $CO_2$ until their confluence reaches about 80%.

BS-C-1 cells (ATCC, Cat#CCL-26) in collagen embedded 3D cultures were immuno-labeled with α-tubulin. 50,000 BS-C-1 cells were centrifuged and re-suspended in 100 μL of 4 mg/mL collagen I (Advanced BioMatrix, Cat#5201-1KIT). The suspension containing collagen I and BS-C-1 cells was then dispensed onto coverslips in 6-well plates. After incubation at 37 °C for 20 min to solidify the collagen, cells on the coverslips were cultured in EMEM (ATCC, Cat#30-2003) with 10% FBS at 37 °C with 5% $CO_2$ until their confluence reaches about 80%.

#### Fixation and labeling of TOM20, α-tubulin, and Nup98

In preparation of immuno-labeling of TOM20 and α-tubulin specimens, cultured cells were first fixed with 37 ºC pre-warmed 3% PFA (Electron Microscopy Sciences, Cat#15710) and 0.5% GA (Electron Microscopy Sciences, Cat#16019) in PBS at RT for 15 min. In preparation of immuno-labeled Nup98 specimens, cultured cells were first rinsed with 37 °C pre-warmed 2.4% PFA in PBS for 20 s, and then extracted with 37 °C pre-warmed 0.4% Triton X-100 (Sigma-Aldrich, Cat#X100) in PBS for 3 min. Then, cells were fixed with 2.4% PFA in PBS for 30 min. After fixation, cells were washed twice with PBS and then quenched with freshly-prepared 0.1% $NaBH_4$ (Sigma-Aldrich, Cat#452882) in PBS for 7 min. Subsequently, cells were washed three times with PBS and then treated with blocking buffer (3% BSA (Jackson ImmunoResearch, Cat#001-000-162) and 0.2% Triton X-100 in PBS for TOM20 and α-tubulin, and 5% BSA in PBS for Nup98) for 1 h, gently rocked at RT. Then, cells were incubated with primary antibodies (Santa Cruz Biotechnology, Cat#sc-11415 for TOM20; Sigma-Aldrich, Cat#T5168 for α-tubulin; Cell Signaling Technology, Cat#2598 for Nup98) at 4 ºC overnight. After washed three times for 5 min each time with wash buffer (0.05% Triton X-100 in PBS), cells were then incubated with secondary antibodies (Molecular Probes, Cat#A21245 and A21236 for Alexa Fluor 647 (referred as 'AF647' hereafter, for simplicity); DNA-conjugated anti-mouse P1, anti-rabbit P1, and anti-rabbit P4 for DNA-PAINT [1–3]) at RT for 5 h. Both primary and secondary antibodies were diluted to 1:500 in antibody dilution buffer (1% BSA and 0.2% Triton X-100 in PBS for TOM20 and α-tubulin, and 5% BSA in PBS for Nup98). After washed three times (5 min each time with wash buffer), cells were post-fixed with 4% PFA in PBS for 10 min. Cells were then washed three times with PBS and stored in PBS at 4 °C until imaging.

**Fixation and labeling of mouse brain sections**

All animal studies were approved by the Indiana University School of Medicine Institutional Animal Care and Use Committee (IACUC). An 8-month-old murine model of Alzheimer's disease (5xFAD) was anesthetized with Tribromoethanol (Avertin) 125−250mg/kg IP and transcardially perfused with saline. Brain was post-fixed with 4% PFA in PBST (0.1% Tween (VWR, Cat#0777) in PBS) for 24 h. Tissue was then transferred to 30% sucrose (Fisher Chemical, Cat#57-50-1). Tissue was embedded in O.C.T. compound (Fisher Healthcare, Cat#23-730-571) and hemibrain was longitudinally sectioned on a cryostat (Leica, Cat#CM1950) at 30 µm thick. Sections were stored at −20 °C in cryoprotectant (30% glycerol (Sigma-Aldrich, Cat#G5516) and 30% ethylene glycol (Sigma-Aldrich, Cat#293237) in PBS). Prior to staining, sections were washed three times in PBST for 10 min each time and then treated for antigen retrieval with 10 mM sodium citrate (Fisher Chemical, Cat#S279-500) and 0.5% Tween in PBST at 85 °C for 10 min. Sections were blocked in normal donkey serum (Sigma-Aldrich, Cat#D9663) for 1 h and incubated with anti-β-Amyloid (Cell Signaling Technology, Cat#2454) at 4 °C overnight. Following three PBST washes, sections were then stained with donkey anti-rabbit AF647-conjugated antibodies (Molecular Probes, Cat#A31573) at RT for 1 h. Both primary and secondary antibodies were diluted to 1:1000 in blocking buffer. Nuclei were stained with DAPI (Sigma-Aldrich, Cat#10236276001) diluted to 1:10,000 in PBST at RT for 2 min. Sections were then wet mounted onto coverslips and dried at 4 °C overnight before imaging.

**Fixation and labeling of mouse forelimb cyrosections**

All murine experiments were approved by the Purdue Animal Care and Use Committee (PACUC). E14.5 mouse embryos were generated by the timed mating of wild type C57Bl/6 mice. Mice were euthanized via $CO_2$ inhalation and confirmed by cervical dislocation. Embryos were removed from the uterine horns and rinsed with PBS. Forelimbs were removed from the embryos and mounted in 1% low gelling agarose (Sigma-Aldrich, Cat#A9045) cubes. Agarose cubes were submerged in 0.05% SDS (VWR, Cat#0837) and 2% Penicillin-Streptomycin in PBS, and gently rocked at RT. The SDS buffer was replaced every 48 h until decellularization was completed (3–5 d). Upon decellularization, agarose cubes were rinsed with PBS for 1 h, and then fixed with 4% PFA (Fisher Scientific, Cat#J19943K2) in PBS for 1 h, rinsed with PBS for 1 h again gently rocked at RT. Forelimbs were removed from the agarose cubes for cryosectioning. Forelimbs were submerged in 15% sucrose (Sigma-Aldrich, Cat#84097) at 4 °C until equilibrated (indicated by the specimen sinking to the bottom of the tube), and then submerged in 30% sucrose at 4 °C until equilibrated. Forelimbs were embedded in O.C.T. compound (Sakura Finetek, Cat#4583), frozen in dry-ice-cooled isopentane, and stored at −80 °C until sectioning. 20-μm-thick cryosections containing cartilage from the humerus were collected on coverslips and stored at −20 °C. Before staining, cryosections were rinsed with PBS for 5 min to remove any residual O.C.T. compound, fixed with 4% PFA in PBS for 15 min, and rinsed with PBS for 5 min again. Cryosections were then quenched with 0.1% $NaBH_4$ in PBS for 15 min, and washed with PBS for 5 min. Cryosections were blocked with 10% donkey serum (Sigma-Aldrich, Cat#S30) and 0.2% BSA (Sigma-Aldrich, Cat#A9418) in PBS for 1 h, and then incubated with AF647-conjugated WGA (Invitrogen, Cat#W32466) diluted to 1:200 in PBS at 4 °C overnight. After that, cryosections were washed three times with PBS and stored in PBS at 4 °C until imaging.

**Imaging buffers and sample mounting**

Immediately before imaging samples labeled with AF647, the coverslip with specimens on top of it was placed on a custom-made holder. 40 μL of imaging buffer (10% (w/v) glucose in 50 mM Tris (Avantor, Cat#JT4109), 50 mM NaCl (Fisher Chemical, Cat#S271-500), 10 mM MEA (Sigma-Aldrich, Cat#M6500), 50 mM BME (Sigma-Aldrich, Cat#M3148), 2 mM COT (Sigma-Aldrich, Cat#138924), 2.5 mM PCA (Sigma-Aldrich, Cat#37580), and 50 nM PCD (Sigma-Aldrich, Cat#P8279), pH 8.0) was added on top of the coverslip. Then another cleaned coverslip was placed on top of the imaging buffer. This coverslip sandwich was sealed with valap. The sample cavity with immuno-labeled cells on the top coverslip was prepared in a similar way by placing the cleaned coverslip at the bottom and the coverslip with cells on top of it with the cell side surface facing down.

Immediately before imaging samples tagged with DNA-PAINT probes, the coverslip with specimens on top of it was placed on a cell chamber (Invitrogen, Cat#A7816). 600 μL of imaging buffer (2 nM ATTO-655 conjugated DNA imager strand diluted in 500 mM NaCl in PBS) was added into the chamber. In the distorted wavefront control experiment, P1 imager strand was used to image TOM20. In Exchange-PAINT imaging, the chamber was mounted firmly on the sample stage to minimize the lateral drift. We first added imaging buffer with P4 strand to image TOM20, and then used syringes to remove the buffer, wash samples with PBS for several times, and add imaging buffer with P1 strand to image α-tubulin.

**Microscope Setup**

The system (Figure S1F) was built around an Olympus IX-73 microscope stand (Olympus America, Cat#IX-73) equipped with a 100×/1.35-NA silicone-oil-immersion objective lens (Olympus America, Cat#FV-U2B714) and a PIFOC objective positioner (Physik Instrumente, Cat#ND72Z2LAQ). Three laser lines at wavelengths of 642 nm (MPB Communications, Cat#2RU-VFL-P-2000-642-B1R), 560 nm (MPB Communications, Cat#2RU-VFL-P-500-560), and 405 nm (Crystalaser, Cat#DL-405-100) were coupled into a polarization-maintaining single-mode fiber (Thorlabs, Cat#PM-S405-XP) after passing through an acousto-optic tunable filter (AA Opto-electronic, Cat#AOTFnC-400.650-TN) for wavelength selection and power modulation. The excitation light coming out of the fiber was focused to the pupil plane of the objective lens after passing through a filter cube holding a quadband dichroic mirror (Semrock, Cat#Di03-R405/488/561/635-t1). The excitation light could be translated sideways by a mirror conjugated to the sample plane for switching between epi-illumination and highly inclined and laminated optical sheet (HILO) imaging modalities [4]. Additionally, a transmitted Köhler illuminator inside the microscope stand equipped with a motorized shutter (Edmund Optics, Cat#87-208) illuminated the sample between acquisition cycles for focus stabilization. Besides, to observe the nucleus labeled with DAPI, an alternative illumination module was used, where light from a mercury light source (Olympus America, Cat#U-LH100HG) was directed by a motorized flip mirror, passed through a filter cube holding a bandpass filter (Chroma, Cat#AT350/50x) and a dichroic mirror (Chroma, Cat#T400LP), and then illuminated the sample. The pupil plane of the objective lens was imaged onto a deformable mirror (Boston Micromachines, Cat#Multi-3.5), which allowed for introducing controlled amount of wavefront aberrations to test the performance of INSPR experimentally. The fluorescent signal was magnified by relay lenses arranged in a $4f$ alignment to a final magnification of ~54, and then was split with a 50/50 non-polarizing beam splitter cube (Thorlabs, Cat#BS016). The separated fluorescent signals were then delivered by two mirrors onto a 90° specialty mirror (Edmund Optics, Cat#47-005), passed through a motorized filter wheel holding five alternative bandpass filters (Semrock, Cat#FF01-731/137-25 and FF01-600/52-25; Chroma, Cat#ET665LP,

ET700/75m, and ET460/50m), and were then projected simultaneously on an sCMOS camera (Hamamatsu, Cat#Orca-Flash4.0v3) with an effective pixel size of 120 nm. The detection planes that received the signals reflected and transmitted by the beam splitter cube were referred as plane 1 and plane 2, respectively (Figure S1G). To adjust the distance between the two detection planes, two piezo inertia actuators (Thorlabs, Cat#PIAK10 and PIA13) were equipped on the mirror that delivered the reflected signal onto the 90° specialty mirror. The imaging system was controlled by a custom-written program in LabVIEW (National Instruments).

#### Definition and characterization of biplane distance

We estimated the distance between two detection planes in the biplane configuration [5] (named as biplane distance $\delta$, Figure S1G) by imaging an isolated fluorescent bead on the coverslip at different axial positions. Intensity profiles (i.e. point spread functions, PSFs) in each z position were fitted with a 2D Gaussian function

$$f(x,y) = I_0 e^{-\left[\frac{(x-x_0)^2+(y-y_0)^2}{2\sigma^2}\right]} + b, \tag{1}$$

where $(x_0, y_0)$ is the center position of each PSF, $I_0$ is the peak intensity, $b$ is the background, and $\sigma$ is the standard deviation representing the width of the PSF. The modulation of the estimated standard deviation $\sigma_i$ in each axial position z was modeled as

$$\sigma_i(z) = \sigma_{i0}\sqrt{1 + \left(\frac{z-c_i}{d_i}\right)^2 + A_i\left(\frac{z-c_i}{d_i}\right)^3 + B_i\left(\frac{z-c_i}{d_i}\right)^4}, \tag{2}$$

where $i$ is the indices of planes ($i = 1,2$), $\sigma_{i0}$ is the standard deviation of PSF when the emitter is at the focal plane, $c_i$ is the position of the focal plane, $d_i$ is the depth of focus, and $A_i$ and $B_i$ are empirical coefficients [6,7]. From the position of each focal plane, we obtained the biplane distance $\delta = |c_1 - c_2|$. When imaging above the coverslip surface, due to the refractive index mismatch, the actual focal position $c_i'$ of each plane is shifted to $c_i n_w / n_o$, where $n_w$ and $n_o$ are the refractive indices of the water-based imaging medium and the immersion medium, respectively [8]. Therefore, the distance between two detection planes in the imaging medium is changed to $\delta n_w / n_o$.

#### Calibration of deformable mirror

The calibration of the deformable mirror (DM) was achieved according to previously described methods [9,10]. Briefly, the response of the membrane surface of the DM can be decomposed into an orthogonal set of mirror deformation modes (DM modes), resembling Zernike-based aberration modes for describing the aberrations in an optical system [11]. We selected 23 DM modes related to Zernike modes from vertical astigmatism to tertiary spherical aberration (Wyant order) for calibration. For each DM mode, we retrieved pupil functions with a phase retrieval algorithm [12,13] for 9 different input amplitudes (–4, –3, –2, –1, 0, 1, 2, 3, 4; unit: λ/2π) by imaging isolated fluorescent beads on the coverslip at 11 different axial positions (in an axial range from –2 to +2 µm with a step size of 400 nm). The amplitude coefficients of 32 Zernike modes (Wyant order, from vertical astigmatism to quaternary spherical aberration) were extracted for different input amplitudes in each DM mode. The amplitude relationship between DM mode and Zernike mode can be described as

$$\Phi_{i,j}^{DM} = \sum_{n=5}^{36} c_{n,j} Z_{n,j} \,, \tag{3}$$

where $\Phi_{i,j}^{DM}$ is the $i$th DM mode with input amplitude $j$ ($j$ = –4, –3, –2, –1, 0, 1, 2, 3, 4), $Z_{n,j}$ and $c_{n,j}$ are the $n$th Zernike mode and its corresponding coefficient with input amplitude $j$ (the first four Zernike modes, piston, x-tilt, y-tilt, and defocus, are removed here). By linearly fitting through the coefficients of Zernike modes as a function of DM mode amplitudes, the relationship between DM mode and Zernike mode can be expressed as

$$\Phi_{i,j}^{DM'} = \sum_{n=5}^{36} c_n' Z_n' \,, \tag{4}$$

where $\Phi_{i,j}^{DM'}$ is the $i$th DM mode, $Z_n'$ and $c_n'$ are the $n$th Zernike mode and its fitting coefficient, respectively. By solving Eq. (4) with least-squares, each Zernike mode can be expressed as a linear combination of DM modes.

We then tested the calibration accuracy for 21 Zernike modes (Wyant order, from vertical astigmatism to tertiary spherical aberration). For each Zernike mode, we retrieved pupil functions for amplitudes at ±1 (unit: λ/2π) by imaging a fluorescent bead on the coverslip at different imaging depths, decomposed the pupil functions into 21 Zernike modes, and obtained their retrieved amplitude coefficients. To eliminate the influence of the aberrations induced by the imaging system itself, we calculated the difference between the retrieved amplitude coefficients from the test Zernike mode at amplitudes of +1 and –1, and divided this difference by 2. After processing 21 Zernike modes, we built a heat map representing the relationship between the input and phase retrieved amplitudes of Zernike modes (Figure S2D). It can be seen that this heat map always gets the largest estimation along the diagonal elements. The average error between the input and phase retrieved amplitude along the diagonal elements is 6% in the first 14 Zernike modes and 27% in the last 7 Zernike modes. Besides, we calculated the root-mean-square error (RMSE) between the 21 phase retrieved amplitudes and the 21 input amplitudes (only the test mode has an amplitude of 1, and others are all 0) for each Zernike mode, and got an average RMSE of 13 mλ for the total 21 test modes.

#### Characterization of sCMOS camera

The characterization of sCMOS camera (the offset, variance, and gain for each pixel on the camera) was estimated according to previously described methods [14,15]. Briefly, the offset $o_q$ for pixel $q$ on the camera was obtained by performing a temporal average on 20,000 frames acquired with the camera's entrance port being covered, as

$$o_q = \frac{1}{M}\sum_{m=1}^{M} s_q^m, \tag{5}$$

where $s_q^m$ is the ADU count at frame $m$ for pixel $q$, and $M$ is the total frame number.

The variance $var_q$ for pixel $q$ was obtained by calculating

$$var_q = \frac{1}{M}\sum_{m=1}^{M} \left(s_q^m\right)^2 - {o_q}^2. \tag{6}$$

To estimate the gain for each pixel, we imaged a fluorescent plastic slide (Chroma, Cat#92001) with the 642-nm laser at different illumination intensities (20,000 frames were recorded in each intensity level). By using the Moore-Penrose pseudo-inverse algorithm, the gain $g_q$ for pixel $q$ can be obtained as

$$g_q = \left(B_q B_q^T\right)^{-1} B_q A_q^T\ , \tag{7}$$

$$A_q = \{\left(v_q^1 - var_q\right), \cdots, \left(v_q^n - var_q\right), \cdots, \left(v_q^N - var_q\right)\}\ , \tag{8}$$

$$B_q = \{\left(D_q^1 - o_q\right), \cdots, \left(D_q^n - o_q\right), \cdots, \left(D_q^N - o_q\right)\}, \tag{9}$$

where $v_q^n$ and $D_q^n$ stand for the temporal variance and average of the ADU counts (20,000 frames) for the $n$th illumination intensity in pixel $q$, respectively, and $N$ is the total number of illumination intensity levels.

#### Data acquisition

The microscope setup is extremely susceptible to sample drift in the axial direction for its long data acquisition time, typically from tens of minutes to hours. To compensate this drift, we implemented a focus stabilization module. Before fluorescence imaging, we recorded a series of bright-field images of the sample along the axial direction (from –1 to +1 μm, with a step size of 100 nm) as reference images. During fluorescence imaging, we recorded a real-time bright-field image of the sample after each acquisition cycle (1000 or 2000 frames, depending on the sample stability), and compared the similarities between this real-time image and reference images by calculating their 2D correlation. The correlation values of the most similar reference image and its 9 adjacent images, together with their z positions, were fitted with third degree polynomials. The z position corresponding to the maximum correlation value in the fitting curve was treated as the sample drift. Then we moved the objective lens in the inverse direction to compensate this drift [16]. In this way, focus stabilization can be achieved during data acquisition.

The single-plane dataset for DM calibration was collected by imaging a fluorescent bead on the coverslip (only acquired in plane 2) over an axial range from –2 to +2 μm with a step size of 400 nm, and taking 10 frames per step with a frame rate of 5 Hz. The amplitude of each DM mode was set to be –4, –3, –2, –1, 0, 1, 2, 3, and 4 (unit: λ/2π).

The biplane datasets for testing the accuracy of DM calibration (Figure S2D), measuring the biplane distance (Figure S1G), and building the *in vitro* model (Figure 3, Figure S3) were separately collected by imaging fluorescent beads on the coverslip over an axial range from –1.5 to +1.5 μm with a step size of 100 nm, and taking 50 frames per step with a frame rate of 10 Hz. The biplane distance was set to 580 nm for distorted wavefront control (Figures 2E and 2F, Movie S2), 286 nm for imaging TOM20 labeled with AF647 (Figure 3, Figure S3, Movie S3), and 558 nm for all the other experiments (Figures 4–6, Figures S4–S6, Movies S4–S7).

The biplane dataset for camera calibration was collected by imaging the fluorescent slide. With the cap covered on the camera's entrance port, 20,000 frames were collected with a frame rate of 50 Hz to calculate the offset and variance of the readout noise for each pixel. With illumination from the 642-nm laser at 9 different intensities (from 20 to 300 photons), 20,000 frames were collected with a frame rate of 50 Hz for each illumination level to estimate the gain for each pixel.

The biological sample was first excited with the 642-nm laser at a low intensity of ~50 W/cm$^2$ to find a region of interest. The depth from this region to the bottom coverslip was measured by recording a first position of the objective lens when the dusts on the bottom coverslip were in focus, then recording a second position of the objective lens when the region of interest was in focus. The difference between these two recorded positions was treated as the depth of this region. Before fluorescence imaging, bright-field images of this region were recorded over an axial range from –1 to +1 μm with a step size of 100 nm as reference images for focus stabilization. Then the blinking data were collected at a laser intensity of 2–6 kW/cm$^2$ and a frame rate of 50 Hz. For distorted wavefront control, 2000 frames were collected for each Zernike-based aberration mode with its amplitude at ±1 (unit: λ/2π). For single-section imaging, 2000 frames were collected per cycle and ~50 cycles were collected. For multi-section imaging, the sample was scanned axially by translating the objective lens with a step size of 400 nm from the bottom to the top of the sample. 1000 or 2000 frames were collected for each cycle in one section, 5–14 sections were collected according to the thickness of the sample, and 9–20 cycles were collected in total (Table S1).

**PSFs generation**

According to scalar diffraction theory [17], the point spread function (PSF) of an imaging system can be calculated from the Fourier transform of the pupil function as

$$\mu_0(x, y, z) = \left|\mathcal{F}[P(k_x, k_y)e^{i2\pi k_z z}]\right|^2, \tag{10}$$

where $\mu_0(x, y, z)$ describes the PSF at position $(x, y, z)$ in the sample space, $\mathcal{F}$ denotes the Fourier transform operator, and $P(k_x, k_y)$ is the pupil function at the back focal plane of the objective lens. The size of the pupil function is limited by $(k_x + k_y)^2 \leq \left(\frac{\mathrm{NA}}{\lambda}\right)^2$, where NA is the numerical aperture of the objective lens and $\lambda$ is the emission wavelength of the emitter. The defocus phase is described by the factor $e^{i2\pi k_z z}$, where $k_z = \sqrt{(\frac{n}{\lambda})^2 - k_x^2 - k_y^2}$, and $n$ is the refractive index of the immersion medium.

The pupil function can be expressed as

$$P\left(k_x, k_y\right) = A\left(k_x, k_y\right) \cdot e^{i\Phi(k_x, k_y)}, \tag{11}$$

where $A\left(k_x, k_y\right)$ and $\Phi\left(k_x, k_y\right)$ are the amplitude and phase of the electric field at the pupil plane, respectively. $\Phi\left(k_x, k_y\right)$ describes the optical aberrations introduced by instrument imperfections and the local biological context, which can be decomposed into a series of Zernike modes [18] as

$$\Phi\left(k_x, k_y\right) = \sum_{n=1}^{N} c_n Z_n(k_x, k_y), \tag{12}$$

where $Z_n(k_x, k_y)$ is the $n$th Zernike mode, $c_n$ is its corresponding coefficient, and $N$ is the number of Zernike modes. In our simulation, $N$ was set to 25. Among these 25 Zernike modes, piston, x-tilt, and y-tilt do not influence the shape of the PSF, and defocus depends on the axial position, so we only considered the rest 21 Zernike modes (Wyant order, from vertical astigmatism to tertiary spherical aberration).

The simulated PSFs in the biplane configuration were generated as follows. (1) Coefficients $(c_0, c_2, \cdots, c_N)$ was set to obtain a pupil function $P\left(k_x, k_y\right)$. (2) Pairs of normalized PSFs $\mu_0(x, y, z)$ and $\mu_0(x, y, z + \delta)$ were generated for two planes on basis of $P\left(k_x, k_y\right)$, where $\delta$ is the biplane distance. Here we assume the two planes share the same pupil function. (3) $\mu_0$ was multiplied with total photon counts $I$ and added with a background count $bg$ to obtain ideal PSFs $\mu = I\mu_0 + bg$. (4) $\mu$ was corrupted with Poisson noise.

In our simulation, the biplane distance $\delta$ was set to 580 nm. The simulation parameters including the emission wavelength, the numerical aperture of the objective lens, the effective pixel size on the detection plane, and the refractive indices of the imaging medium and immersion medium were consistent with our experimental conditions. For simulating the randomly aberrated PSFs (Figures 2A–2D, Movie S1), the amplitudes of 21 Zernike modes were randomly sampled from –1 to +1 (unit: λ/2π) in each trial (30 trials in total). 2000 PSFs with a sub-region size of 40 × 40 pixels were located randomly within an axial range from –800 to +800 nm and a lateral range from –3 to +3 pixels to the center. For each PSF, the total photon counts were sampled from a Gaussian distribution with a mean of 2000 and a standard deviation of 500, and the background count was uniformly sampled from 10 to 20 counts per pixel. For simulating PSFs above the coverslip surface (Figures S2A and S2B), the amplitudes of Zernike modes were set according to our previously estimated optical aberrations (consisting of vertical astigmatism, diagonal astigmatism, vertical coma, horizontal coma and primary and secondary spherical aberrations) at various imaging depths (0, 6.7, 14.35, 27.55, and 45.4 μm) [10]. The axial positions of PSFs were randomly sampled from –800 to +800 nm, but with a defocus offset at a certain depth (Figure S2B). The number of PSFs, sub-region size, lateral range to the center, total photon counts, and background count were the same with those of the randomly aberrated PSFs. For simulation in different signal to background ratio (SBR) conditions (Figure S2C), the amplitudes of 21 Zernike modes were randomly sampled from –1 to +1 (unit: λ/2π) in each trial (11 trials in total). The pairs of the total photon counts and background count were set to (5000, 5), (1000, 5), (5000, 20), (1000, 20), (5000, 100), and (5000, 100), and the number of PSFs was ranging from 100 to 2900 with an increment of 200. The sub-region size, axial range, and lateral range to the center were the same with those of the randomly aberrated PSFs. For simulating the channel-specific PSFs (Figures S2E and S2F), the aberrated PSFs (vertical astigmatism aberration with an amplitude of 2 (unit: λ/2π)) were located at axial positions from –500 to +500 nm, with a step size of 100 nm, and 1000 PSFs were generated for each z position. The total photon counts were 2000, and the background count was 30. Plane 1 and plane 2 were related with an affine transformation with a rotation of 30 degrees. Then, except for considering Poisson noise, the pixel-dependent Gaussian noise was added to each pixel of the simulated datasets (the variance distribution was shown in the inset of Figure S2E).

**INSPR framework**

INSPR constructs an *in situ* 3D PSF response directly from single molecules located in the cellular volume. The key of constructing the 3D PSF is to find the relationship between detected single molecules and their axial positions. Drawing inspiration from $k$-means algorithm [19,20], we assigned $n$ detected single molecules (i.e. PSFs library $X = \{\mathrm{PSF}_1, \mathrm{PSF}_2, \cdots, \mathrm{PSF}_n\}$) into $k$ reference z-stack PSFs (i.e. templates $T = \{\mathrm{PSF}'_1, \mathrm{PSF}'_2, \cdots, \mathrm{PSF}'_k\}, k \ll n$) by calculating

$$\arg\max_{S} \sum_{i=1}^{k} \sum_{j \in S_i} \mathrm{Sim}(\mathrm{PSF}_j, \mathrm{PSF}'_i)\,, \tag{13}$$

where Sim is a function to measure the similarity between detected single molecule $\mathrm{PSF}_j$ and template $\mathrm{PSF}'_i$, and $S_i$ is a set of PSFs that are assigned into the $i$th template group. Our goal is to construct $k$ group sets $S = \{S_1, S_2, \cdots, S_k\}$ so that the PSFs within each group are similar to each other, while the averaged PSF of each group is dissimilar to that of other groups.

To ensure this PSF assignment is unique, we used the biplane configuration to detect $n$ pairs of PSFs from two axially separated image planes $X_{bp} = \{(\mathrm{PSF}_{1,1}, \mathrm{PSF}_{1,2}), (\mathrm{PSF}_{2,1}, \mathrm{PSF}_{2,2}), \cdots, (\mathrm{PSF}_{n,1}, \mathrm{PSF}_{n,2})\}$. INSPR starts with a constant pupil and generates $k$ pairs of templates $T_{bp} = \{(\mathrm{PSF}'_{1,1}, \mathrm{PSF}'_{1,2}), (\mathrm{PSF}'_{2,1}, \mathrm{PSF}'_{2,2}), \cdots, (\mathrm{PSF}'_{k,1}, \mathrm{PSF}'_{k,2})\}$ according to 'PSFs generation' section. All detected PSFs in the library are compared with these templates and then assigned into the most similar template group (assignment step in Figure 1). The process then updates the pupil by using aligned and averaged PSFs from the axially assigned group (update step in Figure 1), and the updated pupil is used to generate new templates. This process usually converges in 6–10 iterations (convergence criteria: the phase difference (measured by RMSE) between two adjacent iterations is smaller than 0.02 λ).

INSPR includes three key components: PSFs library construction, PSFs library assignment, and 3D model estimation (Figure S1A), which are explained as follows.

**PSFs library construction.** PSFs library was constructed from the single molecule dataset, including pairs of emission patterns at random depths in a certain axial range. The raw dataset from two planes was first aligned to the same regions of interest (biplane registration), and then cropped into individual sub-regions containing single emitters (segmentation). This process includes the following two steps.

1. Biplane registration. Images in plane 1 were treated as reference, and images in plane 2 were aligned to plane 1 using affine transformation (including translation, scale, shear, and rotation) [21]. Transformation between two planes can be obtained either by imaging 10–15 fluorescent beads on the coverslip with an axial range from –1 to +1 μm with a step size of 100 nm (50 frames in each step), or by collecting a single molecule blinking dataset (1000 or 2000 frames). The image sequences of beads or blinking dataset in two planes were individually projected into the lateral plane by maximum-intensity projection. Then we calculated the affine matrix based on these projection images in two planes (using 'imregtform' function in MATLAB), and registered the images in plane 2 to plane 1 according to the affine matrix (using 'imwarp' function in MATLAB).

2. Segmentation. After summing the images from registered planes, sub-regions with single molecules were cropped using a segmentation algorithm [14], where two uniform filters with different kernel sizes were used to reduce noise, and a maximum filter was used to find local maximum intensities. The positions of these maximum intensities were localized at the centers of candidate sub-regions. We utilized two intensity thresholds (initial intensity threshold $I_{\text{init}}$ and segmentation threshold $I_{\text{seg}}$) and a distance threshold ($d_{\text{thresh}}$) to make sure that each selected sub-region only contains one molecule with enough brightness. First, the candidate sub-regions were selected if their maximum intensities were higher than $I_{\text{init}}$. Second, the overlapping molecules were rejected if the centers of two sub-regions were closer than $d_{\text{thresh}}$. Third, the rest maximum intensities of sub-regions were filtered out if they were below $I_{\text{seg}}$. For distorted wavefront control (Figures 2E and 2F, Table S2, Movie S2), we set the sub-region size to 40 × 40 pixels, $d_{\text{thresh}}$ to 28, $I_{\text{init}}$ to 30, and $I_{\text{seg}}$ ranging from 35 to 90 in order to get ~5000 single molecules. For other *in situ* model estimations (Figures 3–6, Figures S3–S6, Movies S3–S7), we set the sub-region size to 32 × 32 pixels, $d_{\text{thresh}}$ to 26, $I_{\text{init}}$ to 25, and $I_{\text{seg}}$ to 40. For 3D single molecule localization (Figures 3–6, Figures S3–S6, Movies S3–S7), we set the sub-region size to 16 × 16 pixels, $d_{\text{thresh}}$ to 10, $I_{\text{init}}$ to 25, and $I_{\text{seg}}$ to 40.

In single molecule localization techniques, improving the quality of the detected PSFs will improve the accuracy of *in situ* model estimation and 3D localization. This can be realized with combination of adaptive optics [10,11,22–29], tissue clearing and expansion methods [30–32], and light-sheet illumination approaches [33–35].

**PSFs library assignment.** The detected PSFs in the library were assigned to temporary axial positions following three steps. First, the reference z-stack PSFs in two plane were generated from the initial/estimated pupil function. Second, the detected PSFs were classified into different groups based on their similarities with the reference PSFs. Third, 2D alignment was carried out by calculating the shift distance between detected PSFs and the most similar reference PSFs. The details are as follows.

1. Reference z-stack PSFs (templates) generation. Templates were generated from the initial/estimated pupil function, which was a constant pupil in the first iteration, and iteratively optimized in assignment and update steps. INSPR generated $k$ templates in each plane with an axial range from –1.4 to +1.4 µm ($T_{bp} = \{(\text{PSF}'_{1,1}, \text{PSF}'_{1,2}), (\text{PSF}'_{2,1}, \text{PSF}'_{2,2}), \cdots, (\text{PSF}'_{k,1}, \text{PSF}'_{k,2})\}$), which is sufficient to cover all the detected PSFs. The axial step size of the templates $d_t$ was set to $\delta/2$ or $\delta/4$, where $\delta$ is the biplane distance. The setting of the axial step size makes the templates generated from the pupil function of each plane overlap around the center of the axial range. For simulations (Figures 2A–2D, Figures S2A–S2C, Movie S1) and distorted wavefront control (Figures 2E and 2F, Table S2, Movie S2), we set $\delta$ to

580 nm and $d_t$ to $\delta/4$ (145 nm). For imaging TOM20 labeled with AF647 (Figure 3, Figure S3, Movie S3), we set $\delta$ to 286 nm and $d_t$ to $\delta/2$ (143 nm). For other experiments (Figures 4–6, Figures S4–S6, Movies S4–S7), we set $\delta$ to 558 nm and $d_t$ to $\delta/4$ (139 nm).

2. Classification. Each pair of detected single molecules in PSFs library ($X_{bp} = \{(\mathrm{PSF}_{1,1}, \mathrm{PSF}_{1,2}), (\mathrm{PSF}_{2,1}, \mathrm{PSF}_{2,2}), \cdots, (\mathrm{PSF}_{n,1}, \mathrm{PSF}_{n,2})\}$) was assigned to a certain template group with the highest similarity. The similarity is defined as

$$\mathrm{Sim}(\mathrm{PSF}_j, \mathrm{PSF}'_i) = \frac{1}{2}\left[\mathrm{NCC}(\mathrm{PSF}_{j,1}, \mathrm{PSF}'_{i,1}) + \mathrm{NCC}(\mathrm{PSF}_{j,2}, \mathrm{PSF}'_{i,2})\right], \quad (14)$$

where NCC is the normalized cross correlation between detected PSFs and templates in each plane. The value of NCC varies from –1 to +1, where high NCC represents high similarity between detected PSFs and templates. Therefore, each detected PSF in the library was classified into a certain template group, which formed $k$ group sets $S = \{S_1, S_2, \cdots, S_k\}$. We used similarity threshold $\mathrm{Sim}_{\mathrm{min}}$ and number threshold $N_g$ to select high-similarity PSFs in each group, where PSFs with a similarity lower than $\mathrm{Sim}_{\mathrm{min}}$ or groups with PSFs fewer than $N_g$ were rejected. For simulations (Figures 2A–2D, Figures S2A–S2C, Movie S1), we set $\mathrm{Sim}_{\mathrm{min}}$ to 0.5, and $N_g$ ranging from 5 to 30. For distorted wavefront control (Figures 2E and 2F, Table S2, Movie S2), we set $\mathrm{Sim}_{\mathrm{min}}$ to 0.5 or 0.6, and $N_g$ to 15 or 25. For other experiments (Figures 3–6, Figures S3–S6, Movies S3–S7), we set $\mathrm{Sim}_{\mathrm{min}}$ to 0.6, and $N_g$ to 30.

3. 2D alignment. For each detected PSF, 2D cross correlation was used to calculate the shift distance with its corresponding template. To find the correlation peak from the 2D cross correlation image, Fourier interpolation with 10 times up sampling was used to identify the peak with a sub-pixel size of 12 nm. Then the detected PSF was aligned to its template according to the shift distance. Here two shift modes were used. (1) XY_shift_mode = 'separate shift', meaning that the shift distance was calculated individually for each plane, and the PSFs were aligned to their corresponding templates separately. This mode was used in simulation (Figures 2A–2D, Figures S2A–S2C, Movie S1) and distorted wavefront control (Figures 2E and 2F, Table S2, Movie S2). (2) XY_shift_mode = 'together shift', meaning that the shift distances were calculated together for two planes, and the PSFs of two planes were aligned to the corresponding pair of templates. This mode is more robust especially for data with low SBR, so it was used for experimental data.

Here we used the biplane configuration to avoid degeneracies (Figure S1D). Besides, our framework of INSPR can be also generalized to other configurations, such as using a phase mask to generate non-degenerate PSF shapes [36,37] or using a cylindrical lens with prior knowledge (e.g. positive vertical astigmatism aberrations, Figure S1E).

**3D model estimation.** The classified PSFs in each group were averaged to improve SBR, and then re-arranged by their axial positions. The phase retrieval method [12,13] was carried out to estimate a new pupil function, which described the *in situ* 3D model and was used to generate reference z-stack PSFs in 'PSFs library assignment' section. The details are as follows.

1. Group average. Here $k$ group sets $S = \{S_1, S_2, \cdots, S_k\}$ were formed by PSFs library assignment. In each group, the assigned PSFs were similar to each other and aligned to the center of the template. In order to obtain high-contrast images, these assigned PSFs were first normalized by z-score normalization and then averaged together as

$$\mathrm{PSF}_{i,m}^{\mathrm{Ave}} = \mathrm{Ave}\left(\left\{\mathrm{PSF}_{j,m}: j \in S_i\right\}\right), m = (1,2), \tag{15}$$

   where Ave is the image average operation, $\mathrm{PSF}_{i,m}^{\mathrm{Ave}}$ is the averaged PSF of plane $m$ in group $i$, and $\mathrm{PSF}_{j,m}$ is the normalized PSF of plane $m$ in the library. Thus we obtained $2k$ average images $A_{bp} = \left\{\left(\mathrm{PSF}_{1,1}^{\mathrm{Ave}}, \mathrm{PSF}_{1,2}^{\mathrm{Ave}}\right), \left(\mathrm{PSF}_{2,1}^{\mathrm{Ave}}, \mathrm{PSF}_{2,2}^{\mathrm{Ave}}\right), \cdots, \left(\mathrm{PSF}_{k,1}^{\mathrm{Ave}}, \mathrm{PSF}_{k,2}^{\mathrm{Ave}}\right)\right\}$, and their axial positions $Z_{bp} = \left\{\left(Z_{1,1}, Z_{1,2}\right), \left(Z_{2,1}, Z_{2,2}\right), \cdots, \left(Z_{k,1}, Z_{k,2}\right)\right\}$.
2. PSFs stack re-arrangement. The $2k$ averaged PSFs were re-arranged into an axial range from –1.4 to +1.4 μm. The PSFs with the same axial positions in two planes were merged together. Thus a 3D PSF stack was formed, which is usually from –1 to +1 μm with a step size of 100–200 nm.
3. Pupil generation. The 3D PSF stack was used to generate the *in situ* 3D PSF model by phase retrieval (PR) method [12,13], which is based on Gerchberg-Saxon algorithm and outputs a pupil function to generate the retrieved PSFs within an axial range of ~2 μm. The PR process was carried out with a stack of averaged PSFs, their corresponding axial positions, and system parameters including the numerical aperture of the objective lens, the emission wavelength, the refractive index of the immersion medium, and the pixel size on the detection plane. Here we used two update modes. (1) Z_shift_mode = 'shift'. In this mode, PR was carried out three times iteratively. For each time, PR found x tilt, y tilt, and defocus aberrations from decomposed Zernike modes and compensated these aberrations by shifting the lateral and axial positions of averaged PSFs. We used this mode in simulation, imaging experiments, and distorted wavefront control for low-order aberrations (from vertical astigmatism to secondary diagonal astigmatism except for the spherical aberration). (2) Z_shift_mode = 'no shift'. In this mode the positions were not updated. We used this mode in cases where wavefront distortions were significantly large, since the induced phase wrapping could make Zernike decomposition unreliable. In fact, PR is not the only way to estimate the 3D model in INSPR. Any model-estimation tools developed for single molecule localization, such as feature-based mapping [5,38], interpolation [39,40], and deep learning [41–43], can be utilized to build the 3D model in generalized INSPR.

**3D single-molecule localization using INSPR model transformation**

INSPR models the 3D PSFs through the pupil function (as described in 'PSFs generation' section). For the biplane configuration, the PSFs in each plane can be described as

$$\begin{cases} \mu_1(x,y,z) = I_1 \cdot \mu_0(x,y,z) + bg_1 \\ \mu_2(x,y,z) = I_2 \cdot \mu_0(x,y,z+\delta) + bg_2 \end{cases}, \tag{16}$$

where $\mu_1$ and $\mu_2$ represent the PSF models in two planes, $\mu_0(x,y,z)$ and $\mu_0(x,y,z+\delta)$ are normalized PSFs generated by the pupil function at positions $(x,y,z)$ and $(x,y,z+\delta)$, $\delta$ is the biplane distance, $I_1$ and $I_2$ are the total photon counts, and $bg_1$ and $bg_2$ are the background counts.

The collected data from the sCMOS camera come with statistical properties including Poisson noise and pixel-dependent readout noise [44]. If we directly transform and interpolate data between two detection planes in 3D localization (i.e. treating plane 1 as reference and transforming the data from plane 2 to plane 1, as described in 'INSPR framework' section), the noise distribution will no longer maintain these statistical properties, resulting in imaging artifacts and localization imprecisions (Figures S2E and S2F). To this end, we generated a channel-specific PSF model (i.e., transforming the model instead of transforming the data) for single molecule localization.

First, we carried out segmentation for raw data in plane 2 as follows. (1) The center position $(X_1, Y_1)$ of a cropped sub-region in plane 1 was recorded as described in 'INSPR framework' section. (2) $(X_1, Y_1)$ was transformed by affine transformation to find its corresponding position $(X_2, Y_2)$ in plane 2. (3) A sub-region of raw data in plane 2 was cropped with an integer center $(X_{2\text{int}}, Y_{2\text{int}}) = floor(X_2, Y_2)$, and the non-integer offset was calculated as $(\Delta x, \Delta y) = (X_2, Y_2) - (X_{2\text{int}}, Y_{2\text{int}})$. The noise calibration map (including offset, variance, and gain for each pixel) of the sCMOS camera for each sub-region was cropped in a similar way.

The position relationship of the single molecule between plane 1 and plane 2 (Figure S1B) can be described as

$$(X_{2\text{int}} + x', Y_{2\text{int}} + y', 1) = (X_1 + x, Y_1 + y, 1)\begin{bmatrix} a & b & 0 \\ c & d & 0 \\ e & f & 1 \end{bmatrix}, \tag{17}$$

where $(x,y)$ and $(x',y')$ are the positions of single molecules in the cropped sub-regions of two planes. $\begin{bmatrix} a & b & 0 \\ c & d & 0 \\ e & f & 1 \end{bmatrix}$ is the matrix of affine transformation [21] including six parameters, where $\begin{bmatrix} a & b \\ c & d \end{bmatrix}$ represents scale, shear, and rotation operations, and $(e, f)$ represents the translation operation. Affine transformation is a linear transformation and can be written as

$$\begin{cases} X_{2\text{int}} + x' = a(X_1 + x) + c(Y_1 + y) + e = aX_1 + cY_1 + e + ax + cy \\ Y_{2\text{int}} + y' = b(X_1 + x) + d(Y_1 + y) + f = bX_1 + dY_1 + f + bx + dy \end{cases}. \quad (18)$$

Since the center position of the cropped sub-region was transformed from plane 1 to plane 2 by the same affine matrix

$$\begin{cases} X_2 = aX_1 + cY_1 + e = X_{2\text{int}} + \Delta x \\ Y_2 = bX_1 + dY_1 + f = X_{2\text{int}} + \Delta y \end{cases}, \quad (19)$$

by combining Eqs. (18) and (19), the position relationship between $(x, y)$ and $(x', y')$ in cropped sub-regions can be described as

$$\begin{cases} x' = ax + cy + \Delta x \\ y' = bx + dy + \Delta y \end{cases}, \quad (20)$$

showing that the raw single molecule data and the cropped sub-region share the same scale, shear, and rotation parameters $\begin{bmatrix} a & b \\ c & d \end{bmatrix}$ between two planes, except for the translation parameters $(e, f)$.

Second, to generate the channel-specific PSF model in plane 2, the model should satisfy two conditions. First, it shares the same shape information (scale, shear, and rotation) with the cropped sub-regions. Second, it has the same center position before and after transformation. Therefore, the affine transformation applied to the model $\text{Affine}_c = \begin{bmatrix} a & b & 0 \\ c & d & 0 \\ e' & f' & 1 \end{bmatrix}$ should satisfy

$$(X_c, Y_c, 1) = (X_c, Y_c, 1) \begin{bmatrix} a & b & 0 \\ c & d & 0 \\ e' & f' & 1 \end{bmatrix}, \quad (21)$$

where $(X_c, Y_c)$ is the center position of the cropped sub-region (when performing affine transformation, the upper left corner of the image was defined as the origin, and $X_c$ was equal to half of the sub-region size). The translation parameters $(e', f')$ in $\text{Affine}_c$ can be calculated as $\begin{cases} e' = (1 - a)X_c - cY_c \\ f' = (1 - d)Y_c - bX_c \end{cases}$. Thus, the channel-specific PSF model in plane 2 can be described as

$$\mu_2'(x', y', z) = \text{Trans}\{\text{Affine}_c[\mu_2(0,0,z)], (x', y')\}, \quad (22)$$

where Trans is the translation operation and $\mu_2(0,0,z)$ represents the PSF model of plane 2 at position $(0,0,z)$. The model $\mu_2(0,0,z)$ was first transformed to the channel-specific model $\mu_2'(0,0,z)$ by affine transformation $\text{Affine}_c$, and then translated to the position $(x', y')$ given by Eq. (20).

Third, we directly incorporated the channel-specific PSF model inside the maximum likelihood estimator (MLE) [7,39,45] to estimate seven parameters $(x, y, z, I_1, I_2, bg_1, bg_2)$ by considering the Poisson noise and pixel-dependent sCMOS noise as

$$L(\theta|D) = \prod_q \frac{(\mu_{1q}+\gamma_{1q})^{(D_{1q}+\gamma_{1q})} e^{-(\mu_{1q}+\gamma_{1q})}}{\Gamma(D_{1q}+\gamma_{1q}+1)} \prod_q \frac{(\mu'_{2q}+\gamma_{2q})^{(D_{2q}+\gamma_{2q})} e^{-(\mu'_{2q}+\gamma_{2q})}}{\Gamma(D_{2q}+\gamma_{2q}+1)}, \quad (23)$$
$$\theta \in (x, y, z, I_1, I_2, bg_1, bg_2),$$

where $D$ is the cropped sub-region of two planes, $q$ is the pixel index, $\mu_1$ and $\mu_2{}'$ represent the PSF models in planes 1 and 2, respectively. $\gamma_{mq} = \frac{var_{mq}}{g_{mq}^2}$, where $var_{mq}$ and $g_{mq}$ are the variance and gain for pixel $q$ in plane $m$ ($m$ = 1, 2). $\theta$ denotes the fitting parameters including the same position $(x, y, z)$, and different total photon counts $(I_1, I_2)$ and background counts $(bg_1, bg_2)$ for two planes. A modified Levenberg-Marquadt method [46] was used to optimize $\theta$ by minimizing the negative log-likelihood function

$$-\ln(L) = \sum_q \mu_{1q} - (D_{1q} + \gamma_{1q})\ln(\mu_{1q} + \gamma_{1q}) + \sum_q \mu'_{2q} - (D_{2q} + \gamma_{2q})\ln(\mu'_{2q} + \gamma_{2q}). \quad (24)$$

The first and second derivatives are

$$f = -\frac{\partial \ln(L)}{\partial \theta} = \sum_q \left(1 - \frac{D_{1q} + \gamma_{1q}}{\mu_{1q} + \gamma_{1q}}\right)\frac{\partial \mu_{1q}}{\partial \theta} + \sum_q \left(1 - \frac{D_{2q} + \gamma_{2q}}{\mu'_{2q} + \gamma_{2q}}\right)\frac{\partial \mu'_{2q}}{\partial \theta}, \quad (25)$$

$$f' = \frac{\partial f}{\partial \theta} = \sum_q \frac{D_{1q} + \gamma_{1q}}{\left(\mu_{1q} + \gamma_{1q}\right)^2}\left(\frac{\partial \mu_{1q}}{\partial \theta}\right)^2 + \left(1 - \frac{D_{1q} + \gamma_{1q}}{\mu_{1q} + \gamma_{1q}}\right)\frac{\partial^2 \mu_{1q}}{\partial \theta^2}$$
$$+ \sum_q \frac{D_{2q} + \gamma_{2q}}{\left(\mu'_{2q} + \gamma_{2q}\right)^2}\left(\frac{\partial \mu'_{2q}}{\partial \theta}\right)^2 + \left(1 - \frac{D_{2q} + \gamma_{2q}}{\mu'_{2q} + \gamma_{2q}}\right)\frac{\partial^2 \mu'_{2q}}{\partial \theta^2}, \quad (26)$$

where the second derivatives $\frac{\partial^2 \mu_{1q}}{\partial \theta^2}$ and $\frac{\partial^2 \mu'_{2q}}{\partial \theta^2}$ were set to 0, and the fitting parameters were updated from

$$\theta_{n+1} = \theta_n - \frac{f}{f'(1+\beta)}, \quad (27)$$

where $\beta$ is a damping factor to adjust the convergence speed, and was set to 0 here.

The localization speed in INSPR mainly depends on the speed of fitting the parameters in the 3D PSF model generated by the Fourier transform of the retrieved pupil function. To speed up this process, we used the cubic interpolation methods [47] to pre-generate 3D PSF models along the axial direction for each voxel of 0.25 pixel × 0.25 pixel × 4 nm in the whole range of 25 pixels × 25 pixels × 2.6 μm. The GPU implementation of cubic interpolation achieves a localization speed of 240 PSFs per second, ~400 times faster than the CPU implementation using MATLAB. The code was tested on a computer with an Intel

Core i7-8700K processor at 3.70 GHz with 32 GB memory and an NVIDIA GeForce GTX 1070 graphics card with 8.0 GB memory.

#### Calculation of Cramér-Rao lower bound

To quantify the Fisher information content of detected PSFs in INSPR, the Cramér-Rao lower bound (CRLB) [7] for estimating localization precision in an unbiased estimator was calculated as

$$\mathrm{var}(\theta_i) \geq [F(\theta)^{-1}]_{ii}, \tag{28}$$

where $var(\theta)$ is the estimation variance of an estimator, $F(\theta)$ is the Fisher information matrix, $\theta$ is a vector of estimation parameters, and $i$ denotes the index of each parameter.

By incorporating the noise characteristic (Poisson noise and pixel-independent readout noise) of the sCMOS camera and the channel-specific PSF model, the relevant Fisher information in each element can be calculated as

$$F_{ij}(\theta) = \sum_q \frac{1}{\mu_{1q}+\gamma_{1q}} \frac{\partial \mu_{1q}}{\partial \theta_i} \frac{\partial \mu_{1q}}{\partial \theta_j} + \sum_q \frac{1}{\mu'_{2q}+\gamma_{2q}} \frac{\partial \mu'_{2q}}{\partial \theta_i} \frac{\partial \mu'_{2q}}{\partial \theta_j}, \theta \in (x, y, z, I_1, I_2, bg_1, bg_2), \tag{29}$$

where $\mu_1$ and $\mu'_2$ represent the PSF models in planes 1 and 2, respectively. $\gamma$ is the noise characteristic of the sCMOS camera, and $q$ is the pixel index.

Furthermore, the Fisher information in the x and y dimensions was changed by considering the position relationship in the channel-specific PSF model $\begin{cases} x' = ax + cy + \Delta x \\ y' = bx + dy + \Delta y \end{cases}$ (described by Eq. (20) in '3D single-molecule localization' section), where $(x, y)$ and $(x', y')$ are the positions of the PSF model of two planes, $(\Delta x, \Delta y)$ is the non-integer offset in plane 2, and $\begin{bmatrix} a & b \\ c & d \end{bmatrix}$ represents scale, shear, and rotation operations in affine transformation.

By calculating the derivative of the x dimension in plane 2

$$\frac{\partial \mu'_{2q}}{\partial x} = \frac{\partial \mu'_{2q}}{\partial x'} \frac{\partial x'}{\partial x} + \frac{\partial \mu'_{2q}}{\partial y'} \frac{\partial y'}{\partial x} = a \frac{\partial \mu'_{2q}}{\partial x'} + b \frac{\partial \mu'_{2q}}{\partial y'}, \tag{30}$$

the Fisher information in the x dimension can be written as

$$F_{xx} = \sum_q \frac{1}{\mu_{1q} + \gamma_{1q}} \left( \frac{\partial \mu_{1q}}{\partial x} \right)^2 + \sum_q \frac{1}{\mu'_{2q} + \gamma_{2q}} \left( \frac{\partial \mu'_{2q}}{\partial x} \right)^2 \tag{31}$$

$$= \sum_q \frac{1}{\mu_{1q} + \gamma_{1q}} \left( \frac{\partial \mu_{1q}}{\partial x} \right)^2 + \sum_q \frac{1}{\mu'_{2q} + \gamma_{2q}} \left( \frac{\partial \mu'_{2q}}{\partial x'} \frac{\partial x'}{\partial x} + \frac{\partial \mu'_{2q}}{\partial y'} \frac{\partial y'}{\partial x} \right)^2$$

$$= \sum_q \frac{1}{\mu_{1q} + \gamma_{1q}} \left( \frac{\partial \mu_{1q}}{\partial x} \right)^2 + \sum_q \frac{1}{\mu'_{2q} + \gamma_{2q}} \left( a \frac{\partial \mu'_{2q}}{\partial x'} + b \frac{\partial \mu'_{2q}}{\partial y'} \right)^2 .$$

Similarly, the Fisher information in the y dimension can be written as

$$F_{yy} = \sum_q \frac{1}{\mu_{1q} + \gamma_{1q}} \left( \frac{\partial \mu_{1q}}{\partial y} \right)^2 + \sum_q \frac{1}{\mu'_{2q} + \gamma_{2q}} \left( c \frac{\partial \mu'_{2q}}{\partial x'} + d \frac{\partial \mu'_{2q}}{\partial y'} \right)^2 . \quad (32)$$

**Rejection method**

CRLB is one criterion to measure the localization uncertainty with retrieved 3D PSF model given by the position $(x, y, z)$, photon counts $I$, and background count $bg$. Smaller CRLB means higher localization confidence. Here we focused on the localization uncertainty in the z dimension ($CRLB_z$). In order to improve the quality of reconstructed images, localizations with $CRLB_z$ larger than a certain threshold were rejected. We set this threshold to 30 nm for TOM20 (Figure 3), 40 nm for microtubules in collagen embedded 3D-cultured cells (Figure S6A), 45 nm for low-density plaque Amyloid β (Aβ) (Figure 5A), 50 nm for Nup98 (Figure 4), high-density plaque Aβ (Figure 5K), and Exchange-PAINT imaging (Figure S6E), and 70 nm for cartilage (Figure 6).

Log-likelihood ratio (LLR) is another criterion to measure the similarity between each single molecule dataset and its corresponding PSF model, which can be expressed as

$$\begin{aligned} \text{LLR} &= -2 \ln \left( \frac{L(\mu|D)}{L(D|D)} \right) \\ &= \sum_q 2\left[\mu_{1q} - D_{1q} + (D_{1q} + \gamma_{1q}) \ln(\mu_{1q} + \gamma_{1q}) - (D_{1q} + \gamma_{1q}) \ln(D_{1q} + \gamma_{1q})\right] \\ &+ \sum_q 2\left[\mu'_{2q} - D_{2q} + (D_{2q} + \gamma_{2q}) \ln(\mu'_{2q} + \gamma_{2q}) - (D_{2q} + \gamma_{2q}) \ln(D_{2q} + \gamma_{2q})\right], \end{aligned} \quad (33)$$

where $D$ is the cropped sub-region of single molecule, $\mu$ is the PSF model, $\gamma$ is the noise characteristic of the sCMOS camera, and $q$ is the pixel index. Here we set LLR = 1000 for each 16 × 16 pixels in the single molecule dataset.

Besides, single molecules more than 800 nm out of focus and less than 1000 photon counts were rejected in our reconstructions.

#### Drift correction, optical section alignment, and Exchange-PAINT alignment

To image thick samples, optical sections were recorded as described in 'Data acquisition' section. The drift correction and optical-section alignment were carried out according to a previously described method [10]. In each optical section, drift was calibrated by calculating the correlation between each 3D volume consisting of localized single molecules from 1000 frames using a redundancy-based correction method [9, 48,49]. These calibrated 3D volumes formed an ~1.6-μm-thick optical section. Whole cells or tissue specimens were scanned axially by translating the objective lens with a step size of 400 nm which ensured enough overlapping regions between adjacent optical sections. We performed the 3D correlation based method [9] to align two adjacent optical sections, and finally reconstructed a super-resolution 3D volume.

Exchange-PAINT imaging was performed sequentially, first TOM20 and then α-tubulin, as described in 'Imaging buffers and sample mounting' section. Before fluorescence imaging, we recorded a series of bright-field images of the sample over an axial range from –1 to +1 μm with a step size of 100 nm as reference images for both focus stabilization (as described in 'Data acquisition' section) and Exchange-PAINT alignment. After calibrating the drift in each imaging process, we used these reference images to align the calibrated results from TOM20 to α-tubulin by performing the 3D correlation based method [9].

#### Capability of INSPR in 3D cultured cells and Exchange-PAINT imaging

As an additional demonstration of INSPR, we performed reconstructions of microtubules immuno-labeled with AF647 in collagen embedded 3D-cultured cells, as well as Exchange-PAINT imaging [2] in COS-7 cells (Figure S6). Compared with 2D monolayer cells cultured on flat coverslips, 3D cultured cells can mimick the *in vivo* cell behaviors realistically and provide useful physiological information [50]. We reconstructed a 28.1 μm × 28.1 μm × 4.1 μm volume of microtubules grown in collagen and clearly observed fine and dense microtubules covering the nuclear envelope in the volume (Figures S6A–S6D), showing the capability of INSPR to resolve nanoscale structures in different culture systems. We further utilized INSPR in Exchange-PAINT imaging by sequentially resolving mitochondria and microtubules with high resolution (Figures S6E–S6I), showing the potential capability of INSPR with multiplexed super-resolution labeling approaches.

### Additional details on statistical analysis

#### Simulation analysis

We tested the performance of INSPR in different simulation conditions by generating PSFs with random aberrations (Figures 2A–2D, Movie S1), PSFs at different imaging depths above the coverslip surface (Figures S2A and S2B), PSFs with different SBR conditions (Figure S2C), and channel-specific PSFs (Figures S2E and S2F).

For PSFs with random aberrations, we decomposed the INSPR retrieved pupil into 21 Zernike modes (Wyant order, from vertical astigmatism to tertiary spherical aberration) and recorded their amplitudes. For each trial, the phase error and Zernike amplitude errors were defined as the RMSE between the ground truth and INSPR retrieved pupil. We then calculated their mean and standard deviation in total 30 trials.

For PSFs at different imaging depths above the coverslip surface, the defocus offset (i.e., the axial shift from the actual focal plane) was estimated as follows. First, we generated the 3D PSF in a range from –1.5 to +1.5 μm with a step size of 20 nm. Then, we plotted the PSF intensity curve at the center of the lateral position along the axial direction and recorded the axial position of the maximum intensity. After that, we calculated the axial offset between the plane of the maximum intensity and the actual focal plane. In the actual imaging process, we only collected single molecules at the axial plane with the maximum intensity and treated this plane as the focal plane. To simulate this process, we generated 2000 PSFs located randomly with an axial range from –800 to +800 nm, but with a certain defocus offset at each imaging depth, and retrieved *in situ* pupils using INSPR. For each imaging depth, we first generated the 3D PSF by our previously measured optical aberrations [8] with an axial range from –800 to +800 nm and a step size of 100 nm as reference (ground truth). Then we calculated the 3D normalized cross correlations (NCCs) between the reference PSFs and PSFs generated by the Gaussian model, theoretical index mismatch model [9], and INSPR model for different axial offsets. Finally, we recorded the maximum NCC values and their corresponding 3D PSF at different imaging depths.

For different SBR conditions, the amplitude error of Zernike modes was calculated at different photon and background conditions (11 trials in each condition). In each condition, we calculated the mean and standard deviation of the amplitude error in the total 11 trials for different numbers of single molecules. After calculating all conditions, we obtained the convergence curves.

For channel-specific PSFs, the CRLBs in the x, y, and z dimensions were calculated as described in 'Calculation of Cramér-Rao lower bound' section. In the measurement system, the achievable estimation precision is limited by CRLB for an unbiased estimator. The precision in each dimension at a certain position (ranging from –500 to +500 nm with a step size of 100 nm) was calculated from the standard deviation of the estimated positions of 1000 PSFs. The bias in each dimension was calculated as the difference between the averaged position of 1000 PSFs and the ground truth position.

**Distorted wavefront verification**

We tested the accuracy of distorted wavefront estimation from single molecule blinking dataset (Figures 2E and 2F, Movie S2). The deformable mirror was calibrated to introduce individual Zernike-based aberration modes (Wyant order, from vertical astigmatism to tertiary spherical aberration, 21 modes in total) as described in 'Calibration of deformable mirror' section. For each Zernike modes, the single molecule emission patterns were distorted by the introduced Zernike-based aberrations (amplitudes at ±1, unit: $\lambda/2\pi$) and fed into INSPR after acquisition to retrieve the *in situ* PSF and its corresponding pupil function (parameters used for INSPR are shown in Table S2). We decomposed the retrieved pupil functions into 21 Zernike modes, and obtained their amplitude coefficients. To eliminate the influence of instrument imperfections which introduces an offset to the retrieved amplitude, we calculated the difference between the retrieved amplitude coefficients when the amplitude of the input Zernike mode equals to +1 and –1, and divided this difference by 2. After processing 21 Zernike modes, we built a heat map representing the relationship between the input and output amplitudes of Zernike modes. The estimation error between using INSPR and using the *in vitro* model for each Zernike mode was carried out by calculating the difference of the amplitudes at the diagonal elements of the heat map. Besides, we calculated the RMSE between the 21 *in situ* retrieved amplitudes and the 21 *in vitro* retrieved amplitudes for each Zernike mode, and got an average RMSE of 23 mλ for the total 21 test modes.

**Profile quantification of 3D super-resolution reconstructions**

The intensity profiles in reconstruction images were measured with the line profile tool in ImageJ, and then fitted with Gaussian functions. To obtain reliable profiles, we used a pixel size of 12 nm and a standard deviation of 2 pixels (Gaussian blur) in reconstruction images. For TOM20 (Figure 3, Figure S3, Movie S3), we generated reconstruction images of 25 outer membrane structures in the y'-z plane, and obtained their intensity profiles along both y' and z directions. Here the orientation of the cross section was rotated to allow projection of the 3D membrane bounded structures to the 2D image. For Nup98 (Figure 4, Figure S4, Movie S4), we got intensity profiles of 40 Nup98 structures in the x-y plane and 20 Nup98 structures in the x-z plane. For Aβ (Figure 5, Figure S6, Movies S5 and S6), we got intensity profiles of 40 fibrils in both x-y and x-z planes. For cartilage (Figure 6, Figure S6, Movie S7), we got intensity profiles of 15 long elastic fibers (3–5 measurements for each, 53 measurements in total) and 7 short elastic fibers (single measurement for each) in the x-y plane and 40 fibers in the x-z plane. Intensity profiles of TOM20 in the y'-z plane (along the y' or z direction) and Nup98 in the x-y plane (along the y direction) were fitted with a linear combination of two Gaussian functions $f(x) = a_1 e^{-\frac{(x-\mu_1)^2}{2{\sigma_1}^2}} + a_2 e^{-\frac{(x-\mu_2)^2}{2{\sigma_2}^2}} + b$, where $(a_1, a_2, \mu_1, \mu_2, \sigma_1, \sigma_2, b)$ are fitting parameters, $\mu_1$ and $\mu_2$ are the positions of the centers of the peaks, $\sigma_1$ and $\sigma_2$ are the standard deviations representing the widths of structures. Intensity profiles of Nup98 in the x-z plane (along the z direction), and Aβ and cartilage in both x-y and x-z planes (along the direction perpendicular to the fibers) were fitted with a Gaussian function $f(x) = ae^{-\frac{(x-\mu)^2}{2\sigma^2}} + b$, where $(a, \mu, \sigma, b)$ are fitting parameters, $\mu$ is the position of the center of the peak, $\sigma$ is the standard deviation. The full width at half maximum (FWHM) of the Gaussian function equals to $2\sqrt{2\ln 2}\sigma \approx 2.355\sigma$.

For TOM20 (Figure 3, Figure S3), the 3D PSFs in x-y and x-z views were generated by the retrieved pupil functions (as described in 'PSFs generation' section). We first generated a 3D PSF by the *in vitro* model (retrieved from fluorescent beads attached to the coverslip surface) with an axial range from –800 to +800 nm with a step size of 100 nm as reference. Then, in each optical section, we generated a series of *in situ* PSFs in the same axial range, but with a defocus offset (away from the focus plane from –500 to +500 nm with a step size of 50 nm). By calculating the 3D cross correlation, we found the best matched 3D PSF and assigned this 3D PSF to a given optical section. The amplitude of retrieved pupil function $A(k_x, k_y)$ in each optical section was normalized such that $\iint \left|A(k_x, k_y)\right|^2 dk_x\, dk_y$ equals to 1, and the size of the pupil function is limited by $(k_x + k_y)^2 \le (\frac{\text{NA}}{\lambda})^2$, where NA is the numerical aperture of the objective lens and $\lambda$ is the emission wavelength of the emitter.

**Data and software availability**

The INSPR toolbox for *in situ* model estimation and 3D localization is available upon request from the corresponding author and also as supplementary software after publication. Further updates will be made freely available at https://github.com/HuanglabPurdue/INSPR. The software package features an easy-to-use user interface including all steps of 3D single-molecule localization from INSPR model generation, pupil based 3D localization (running on GPU with cubic spline implementation), drift correction, volume alignment, to super-resolution image reconstruction. Additional datasets are available from the corresponding authors upon request.
